# Physicochemical balancing facilitates fast cyclic immunolabeling in centimeter-scale FFPE tissues

**DOI:** 10.64898/2026.03.09.710440

**Authors:** Ya-Hui Lin, Chao-Yin Huang, Yin-Hsu Chen, Yen-Hui Chen, Ping-Liang Ko, Heng-Hua Hsu, Yi-Chung Tung, Yu-Fang Chen, Hui-Chi Chen, Ann-Shyn Chiang, Chih-Ming Ho, Da-Jeng Yao, Kimberly L. Fiock, Kuo-Chuan Wang, Chin-Hsien Lin, Shang-Hsiu Hu, Bi-Chang Chen, Li-An Chu

**Affiliations:** Department of Biomedical Engineering and Environmental Sciences, National Tsing Hua University, Hsinchu, 300044, Taiwan; Brain Research Center, National Tsing Hua University, Hsinchu, 300044, Taiwan; Institute of Information Systems and Applications, National Tsing Hua University, Hsinchu, 300044, Taiwan; Institute of Biomedical Sciences, Academia Sinica, Taipei, 115024, Taiwan; Research Center for Applied Sciences, Academia Sinica, Taipei, 115024, Taiwan; Department of Physics, Fu Jen Catholic University, New Taipei, 242062, Taiwan; Institute of Systems Neuroscience, National Tsing Hua University, Hsinchu, 300044, Taiwan; Department of Mechanical and Aerospace Engineering University of California, Los Angeles, Los Angeles, California 90095, United States; Department of Power Mechanical Engineering National Tsing Hua University, Hsinchu, 300044, Taiwan; Iowa Neuropathology Resource Laboratory, University of Iowa, Iowa City, Iowa 52242, United States; Department of Neurosurgery, National Taiwan University Hospital, Taipei, 100225, Taiwan; Department of Biomedical Engineering, National Taiwan University, Taipei, 106319, Taiwan

**Keywords:** tissue clearing, multiplexed staining, FFPE, archived tissues, AD, CSR, whole-mouse brain imaging

## Abstract

Archived formalin-fixed, paraffin-embedded (FFPE) tissues, including brain-bank specimens, are valuable resources for disease research, but deep and rapid cyclic immunolabeling of intact FFPE specimens remains challenging. Here, we present FIDELITY, an epoxy-free workflow that balances optical transparency, immunostaining performance, and tissue integrity for multi-cycle protein profiling in centimeter-thick FFPE specimens. Neural network-guided optimization identified an SDS-glycine formulation that integrates delipidation and epitope retrieval while preserving tissue stiffness, replacing prolonged epoxy reinforcement and subsequent high-temperature antigen retrieval with a single aqueous pretreatment. Application to human Alzheimer’s disease specimens enabled volumetric imaging and quantitative analysis while retaining compatibility with conventional histopathology. Its compatibility with active electrophoretic staining further supported rapid labeling and five rounds of immunolabeling in a whole FFPE mouse brain. Two-round labeling of a human glioma specimen further demonstrated the applicability of the cyclic workflow to human pathological tissue. Together, FIDELITY provides a practical platform for large-volume, multiplexed protein profiling of intact FFPE specimens.

## Introduction

Formalin-fixed, paraffin-embedded (FFPE) specimens constitute an invaluable resource for human disease research because of their common use for routine preservation in pathology archives and association with clinicopathological records ^1–3^. A large number of multiplexed antibody-based imaging methods have been developed for thin (2-10 µm) FFPE tissue sections, providing proteomic and transcriptomic information ^4–6^. Recently published 3D CyCIF performed high-round multiplexing in 30–50-µm-thick FFPE sections to completely cover the entire tumor cells and present lossless three-dimensional (3D) cell phenotyping; however, the relatively small fraction of the tissue fails to reveal continuous neuron projections and track structural changes across brain regions ^7^.

The organ-wise tissue clearing is critical to visualize neuronal system. Over the last decade, emerging tissue clearing methods, combined with deep immunolabeling techniques and optical sectioning microscopy, have enabled 3D visualization of biomolecules at the centimeter scale, opening up new horizons in basic neuroscience research ^8–12^. Morphology-preserving approaches such as SeeDB and SOLID have enabled large-scale reconstruction of neuronal architecture in fixed brain tissue ^13,14^. For FFPE specimens, some of aqueous tissue-clearing methods, including CLARITY ^15^ and CUBIC ^16,17^ have been adapted and achieved up to 500 µm clearing and staining depth ^18,19^. Protocols derived from iDISCO ^20^, the solvent-based tissue clearing method integrated with immunostaining procedure have much better performance: DIPCO achieved 1.6 mm staining thickness of FFPE lymphatic tumors ^21,22^, and aDISCO ^23^ successfully clear and label various FFPE human specimens of around 5 mm thickness.

These applications have demonstrated the feasibility and value of 3D FFPE pathology, however, 3D cyclic immunostaining of centimeter-scale FFPE specimens remains challenging.

The major challenge in 3D cyclic immunostaining of large FFPE specimens lies in balancing staining efficiency, tissue integrity and epitope retrieval. In HIF-Clear protocol we published in 2024, we adapted electrophoretic immunostaining ^24^ to achieve fast deep labeling, while balancing tissue integrity and epitope retrieval via SHIELD epoxy stabilization ^25^ and heat-induced epitope retrieval (HIER) under extreme high temperature and pressure ^26^. HIF-Clear enables fast immunostaining of a whole mouse brain (from 8 mm to 1 cm) within one day and supported six rounds of cyclic immunostaining, however, the staining depth in human specimen is shallower than it is in mouse tissue ^26^, indicating a more efficient epitope retrieval in need. Moreover, the epoxy stabilization requires five additional days for processing, and ineffective epoxy treatment could result in tissue deformation during electrophoretic processing (Extended Data Figure 1). Together, replacing the epoxy treatment and extreme HIER with a new technique being compatible with electrophoretic staining, retaining tissue integrity and molecular access of recovered epitopes, became a major objective for further workflow development.

In this study, we developed FIDELITY (<u>F</u>FPE-based <u>I</u>terative immunolabeling via <u>D</u>elipidation and <u>E</u>pitope retrieval for <u>L</u>arge-volume Imaging while preserving <u>T</u>issue integrit<u>y</u>), an epoxy-free tissue-processing technique that balanced physical strength required for active electrophoretic staining and chemical delipidation and epitope availability of FFPE specimens. FIDELITY enables volumetric labeling and imaging of archived human Alzheimer’s disease and glioma specimens, while retaining compatibility with routine histological examination. Also, the tissue mechanical strength enables five rounds of active electrophoretic staining and further atlas registration on a mouse brain. Collectively, combined with electrophoretic staining, FIDELITY provides a fast and robust platform for iterative molecular mapping of large archived FFPE specimens.

## Results

### Stepwise screening and neural network–based CSR optimization identify the FIDELITY formulation

HIER is an essential step for recovering epitopes masked by formalin fixation and is therefore critical for immunohistochemical analysis of FFPE specimens ^27–29^. Detergent-containing HIER methods can additionally remove residual lipids from FFPE specimens, thereby increasing optical transparency and facilitating 3D visualization ^26^. However, detergents can disrupt membrane structures and potentially compromise tissue integrity, which is critical to efficient cyclic staining and subsequent cross-round dataset registration. Therefore, optical transparency, immunostaining performance, and tissue integrity represent three interdependent criteria that must be carefully balanced (Figure 1A).

**Figure 1.**
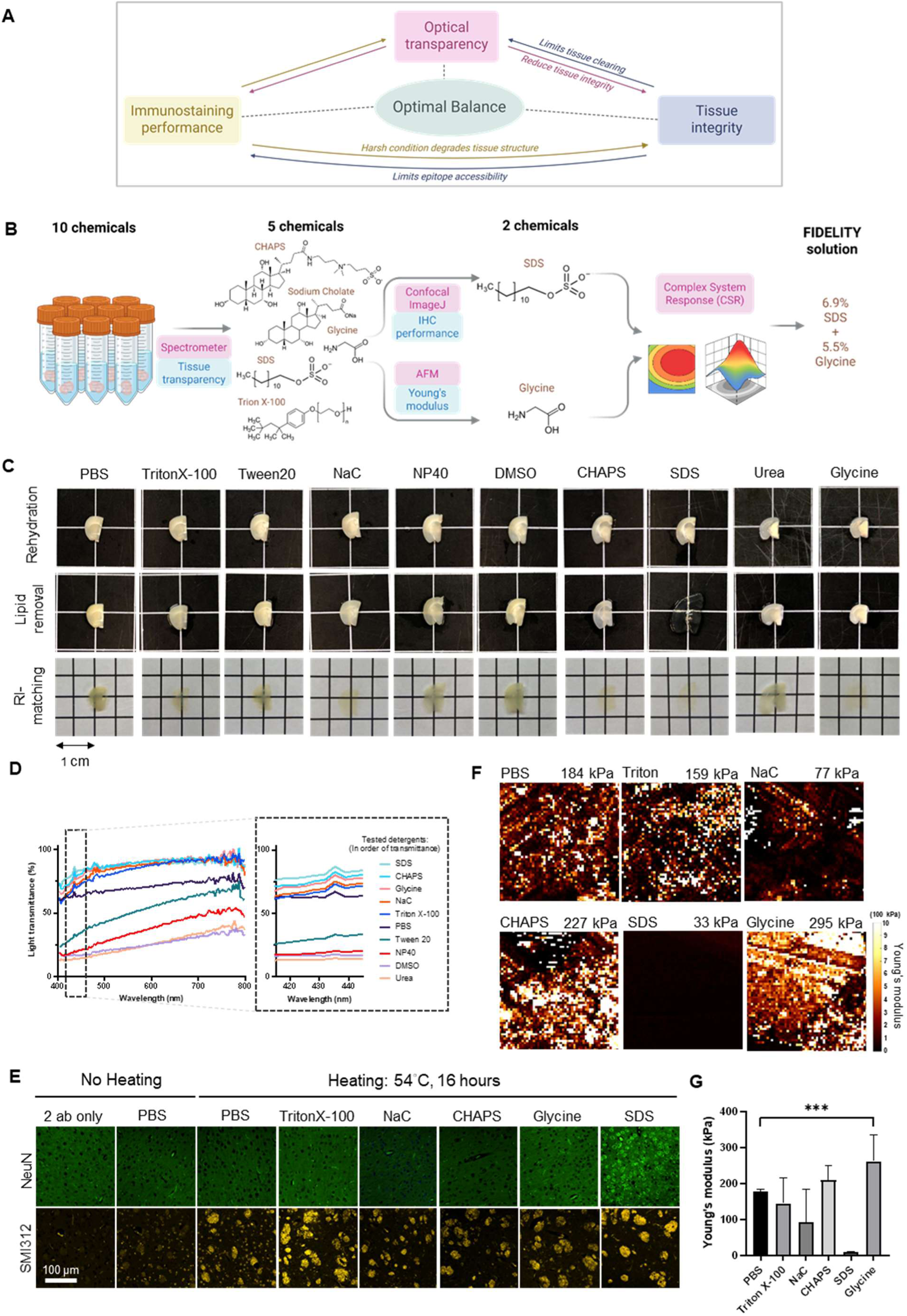
Development of FIDELITY solution. (A) Trade-offs among optical transparency, immunostaining performance, and tissue integrity. Optimization of FFPE tissue processing requires balancing these three interdependent criteria. (Created in https://BioRender.com) (B) Experimental overview of the screening process for FIDELITY chemicals. (Created in https://BioRender.com) (C) 2-mm-thick slices of FFPE mouse brain treated with various detergents or chaotropes. Gross views of the brain slices after rehydration, delipidation, and RI matching are shown. (D) Transparency of the RI-matched brain slices shown in panel C. Detergents that resulted in higher transparency than PBS were selected. (E) Multi-point confocal fluorescence images of neuronal nuclei in the cortex (top row) and axons in the striatum (bottom row) from 10-µm-thick FFPE mouse brain sections treated with detergents selected based on the transparency measurement. (F) Representative heatmaps showing Young’s modulus values according to the corresponding color scale. Young’s modulus was measured in 2-mm-thick FFPE mouse brain slices treated with the detergents identified in panel C. The scan area was 10 µm × 10 µm, with a resolution of 64 × 64 pixels. The average elastic modulus is indicated above each representative map. (G) Comparison of measured Young’s modulus in 2-mm-thick FFPE mouse brain slices treated with the detergents identified selected based on the transparency measurement. Statistical analysis: n = 3 ROIs per group. Data represent mean ± SD. ns: not significant (P > 0.05), *p < 0.05, **p < 0.01, ***p < 0.001, ****p < 0.0001, one-way ANOVA with Tukey’s multiple comparison test.

We sought to develop a single aqueous treatment that could efficiently retrieve antigen epitopes and remove residual lipids while retaining the tissue rigidity required for active immunolabeling. To identify suitable chemicals for the new step, we fixed the incubation conditions at 54 °C for 16 hours (based on the FLASH protocol) ^30^ and screened nine candidate reagents frequently reported to remove lipid or enhance staining performance: Tween 20, Triton X-100, and NP40 (non-ionic detergents); SDS and sodium cholate ^31^ (anionic detergents); CHAPS ^32^ (zwitterionic detergent); DMSO and urea (polar organic agents); and glycine, a small molecule that quenches fixatives and assists antibody elution ^33–35^. Each reagent was prepared as a 10% (w/w) aqueous solution and evaluated for its effects on optical transmittance, tissue stiffness, and immunostaining performance in FFPE mouse brain specimens.

We designed a stepwise screening to narrow down the candidate components (Figure 1B). First, we measured optical transmittance using a custom-built spectrometer and identified five chemicals (Triton X-100, SDS, CHAPS, sodium cholate, and glycine) that improved transmittance relative to PBS controls (Figure 1C and D). We next evaluated their effects on immunostaining using antibodies against NeuN and SMI312. Although SMI312 labeling was detectable across all treatment groups, NeuN labeling was detectable only after SDS treatment (Figure 1E). In parallel, we assessed tissue stiffness by measuring Young’s modulus using atomic force microscopy (AFM) ^36^. Among the same five chemicals, only glycine significantly increased tissue stiffness relative to the PBS-treated control group (Figure 1F and G).

Together, these results nominated SDS as a component with efficient epitope recovery and glycine as a component associated with increased tissue stiffness. SDS alone supported robust antigen retrieval; however, this effect was markedly attenuated in the presence of 10% glycine (Extended Data Figure 2). Interestingly, although glycine alone did not recover NeuN immunoreactivity, it enhanced SDS-mediated immunostaining within a specific concentration range (Figure 2A). These findings indicated a concentration-dependent, non-additive interaction between SDS and glycine.

**Figure 2.**
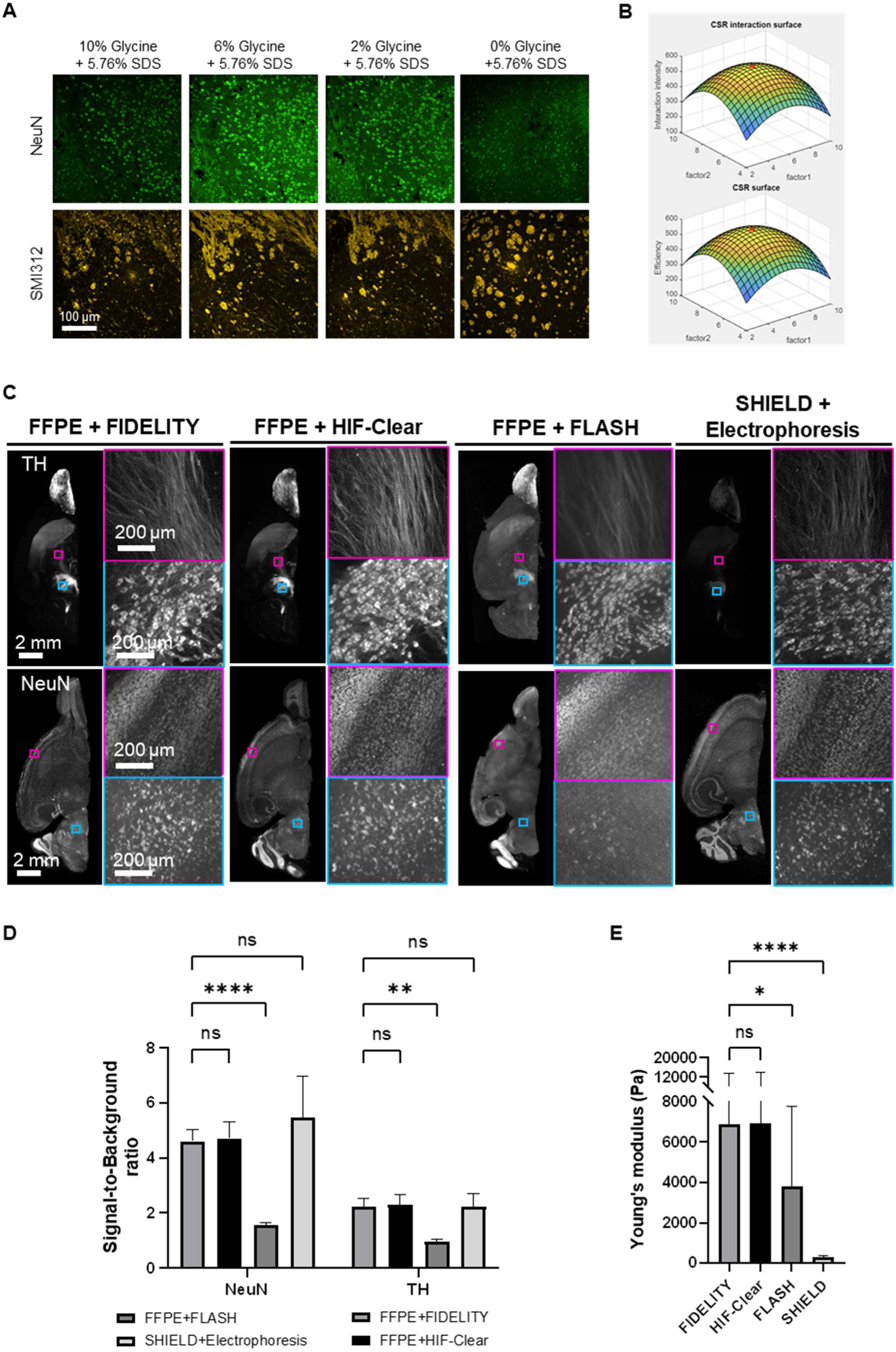
FIDELITY balance the sample robustness, permeability to achieve highest labeling efficiency across whole mouse brain. (A) Multi-point confocal fluorescence images of neuronal nuclei in the cortex (top row) and axons in the striatum (bottom row) from 10-µm-thick FFPE mouse brain sections treated with various ratios of SDS and glycine. (B) Interaction surface generated by Complex Surface Response (CSR) analysis. The peak of the surface (red dot) indicates the predicted optimal condition. (C) Mouse brain hemispheres treated with FIDELITY and other established protocols. The specimens were stained with tyrosine hydroxylase (TH) and neuronal nuclei (NeuN) antibodies and imaged using light-sheet microscopy. (D) Quantitative analysis of immunolabeling performance across FIDELITY and other established protocols. (E) Comparison of measured Young’s modulus in 500-µm-thick slices of mouse brain tissue treated with different tissue-clearing protocols: HIF-Clear, FIDELITY, FLASH, and SHIELD. Statistical analysis: (D) (E) n = 120 positions per group; data represent mean ± SD. ns: not significant (P > 0.05), *p < 0.05, **p < 0.01, ***p < 0.001, ****p < 0.0001, one-way ANOVA with Tukey’s multiple comparison test.

To identify the optimal formulation within this interacting concentration space, we employed a neural neural network–based Complex System Response (CSR) framework ^37,38^. Calibration experiments were selected using an Orthogonal-Array-Composite-Design (OACD) ^39^, and the resulting data were used to fit the response surface. Using NeuN signal-to-background ratio as the optimization output, the model predicted a formulation containing 6.9% SDS and 5.5% glycine (Figure 2B).

We next compared the CSR-predicted formulation with existing protocols. The SDS/glycine solution produced a NeuN signal-to-background ratio that was not significantly different from those obtained with HIF-Clear and SHIELD and was significantly higher than that obtained with FLASH (Figure 2C and D). The Young’s modulus of SDS/glycine-treated tissue was significantly higher than that of SHIELD-treated tissue and was similar in magnitude to that of HIF-Clear-treated tissue, despite the absence of epoxy fixation (Figure 2E). Eliminating epoxy post-fixation also shortened the whole-mouse-brain processing workflow to four days, representing an approximately 50% reduction relative to HIF-Clear and an approximately 65% reduction relative to SHIELD (Figure 3A). We therefore termed this method FIDELITY, an acronym for FFPE-based Iterative immunolabeling via Delipidation and Epitope retrieval for Large-volume Imaging while preserving Tissue integritY.

**Figure 3.**
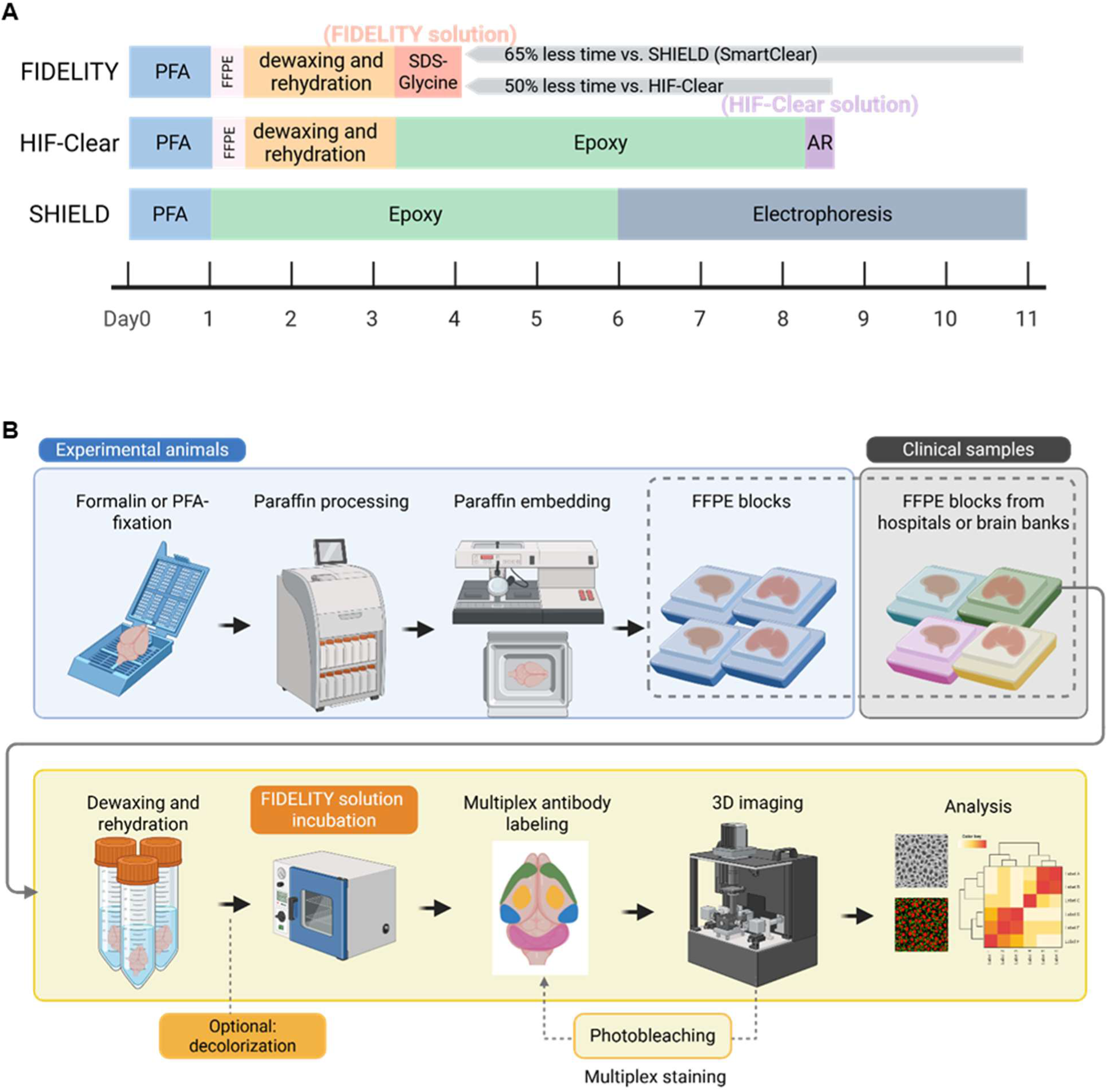
Timeline comparison and workflow of FIDELITY for the following active cyclic immunolabeling of FFPE brain tissues. (A) Comparison of the processing timelines for FIDELITY, HIF-Clear, and SHIELD in preparing whole mouse brains for immunostaining. For SHIELD, the timeline shown is based on the workflow using the SHIELD kit with SmartClear II. SHIELD requires ∼11 days, HIF-Clear requires 8–9 days, and FIDELITY reduces the preparation time to 4 days. (Created in https://BioRender.com) (B) Schematic overview of the FIDELITY pipeline. Brain tissues from experimental animals are fixed with formalin or PFA, processed, and embedded in paraffin to generate FFPE blocks. Archived clinical FFPE brain samples from hospitals or brain banks can also be used. FFPE samples are dewaxed and rehydrated before delipidation and antigen retrieval with the FIDELITY solution, followed by multiplex antibody labeling, light-sheet imaging, and data analysis. Fluorescence signals can be removed by photobleaching to enable iterative rounds of staining and imaging. (Created in https://BioRender.com)

### FIDELITY establishes a streamlined epoxy-free workflow for centimeter-scale FFPE specimens

Details of the FIDELITY pipeline are described in the Materials and Methods section. Briefly, specimens can be either archived FFPE blocks or newly collected samples, as illustrated in Figure 3B.

Newly collected specimens should be thoroughly fixed in 4% PFA or 10% neutral-buffered formalin for 2–7 days prior to paraffin embedding. The resulting FFPE blocks can be stored stably at room temperature for extended periods.

Specimens are firstly processed through dewaxing and rehydration steps (see Materials and Methods for details). Pigment-rich tissues, including non-perfused specimens (containing heme), skin, retina, certain tumors (containing melanin), and samples from aged animals (which may contain lipofuscin), require decolorization with 10% H₂O₂. Decolorization time varies depending on sample properties and conditions, such as tissue size, fixation status, and pigment deposition. For large human specimens, the decolorization process may take 2–3 days (Extended Data Figure 3).

Specimens are then incubated in FIDELITY solution (6.9% SDS and 5.5% glycine in 200 mM borate buffer) at 54 °C for 16 hours. This single aqueous treatment combines delipidation and epitope retrieval and replaces the prolonged epoxy post-fixation used in HIF-Clear. After washing, the specimens are ready for immunostaining. Together with the mechanical measurements shown in Figure 2E, this workflow established a streamlined epoxy-free processing route that retained the tissue stiffness required for subsequent electrophoretic labeling experiments. For freshly collected animal specimens, tissue transparency was also maintained when the paraffin embedding step was omitted (Extended Data Figure 4). This finding suggests that the FIDELITY workflow could be further shortened for specimens that do not require long-term storage.

### FIDELITY enables volumetric labeling of archived human AD specimens while preserving histological morphology

To evaluate the applicability of FIDELITY to archived human FFPE tissue, we used blocks from two clinically and neuropathologically confirmed Alzheimer’s disease (AD) cases and one control case obtained from the Iowa Neuropathology Resource Laboratory. According to the National Institute on Aging–Alzheimer’s Association ABC criteria ^40^, the two AD specimens were classified as A3, B3, C3 (Sample 1) and A3, B3, C2 (Sample 2), where the A, B, and C scores summarize β-amyloid plaque distribution, neurofibrillary tangle stage, and neuritic plaque density, respectively. FIDELITY rendered the entire 5–6-mm-thick human amygdala blocks optically transparent and enabled whole-tissue immunolabeling (Figure 4A and 4C, Extended Data Figure 3 and 5, Supplementary video 1). To assess histological preservation after FIDELITY processing, the FIDELITY-processed tissues were re-embedded in paraffin and sectioned for hematoxylin and eosin (H&E) staining. Comparisons of sections obtained before and after FIDELITY showed preserved cytoarchitecture, maintained vessel-wall organization, and distinguishable neuronal and glial morphologies at the microscopic level (Figure 4B). These results prove that FIDELITY can successfully be applied to archived FFPE brain blocks for volumetric immunostaining while retaining compatibility with conventional histopathological examination.

**Figure 4.**
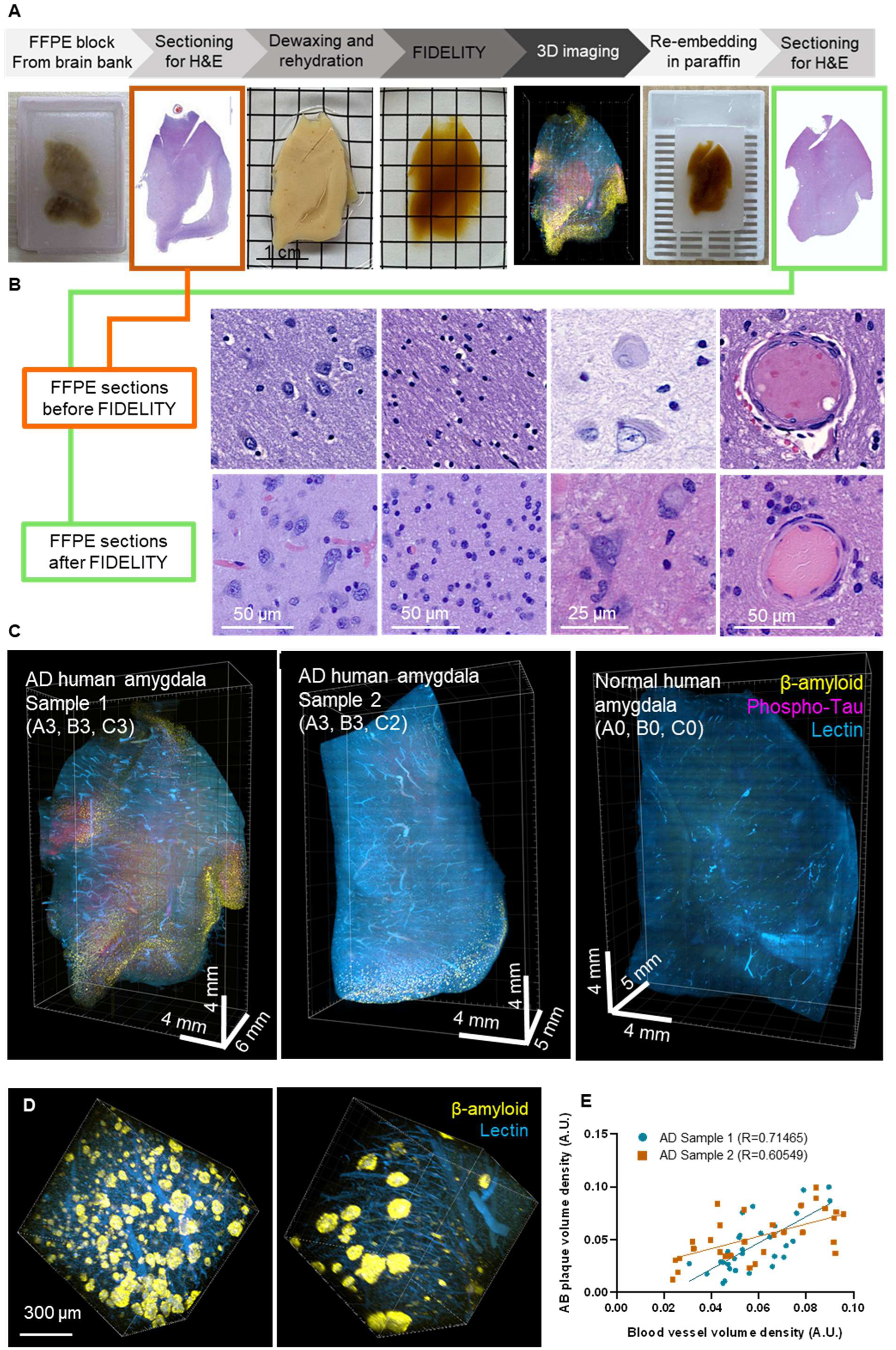
Application of FIDELITY to archived FFPE brain tissues from Alzheimer’s disease patients. (A) Workflow of FIDELITY applied to an FFPE block, followed by re-embedding after 3D imaging. (B) Hematoxylin and eosin (H&E) staining of FFPE sections sectioned from the original human amygdala block (before FIDELITY) and the re-embedded block (after FIDELITY). Representative features, including neuronal cells, glial cells, ballooned neurons, and blood vessels, are indicated from left to right. (C) 3D renderings of FFPE human amygdala specimens. (D) 3D renderings of ROIs with varying levels of Aβ plaques. ROI size = 1 mm³. (E) Correlation analysis of Aβ plaque volume density and blood vessel volume density in two AD specimens. AD Sample 1 (CERAD C = 3): R = 0.71, P < 0.0001; AD Sample 2 (CERAD C = 2): R = 0.61, P = 0.0004.

### Volumetric analysis reveals local associations between Aβ plaque density and vascular density in AD amygdala

Because Aβ pathology and cerebrovascular alterations have been reported closely linked in AD ^41–43^, we used the volumetric datasets to examine local spatial relationships between Aβ plaques and vascular structures. Interestingly, we observed that regions with dense Aβ plaque deposition were frequently accompanied by increased vascular structures (Figure 4D, Supplementary video 2). To quantify this relationship within individual specimens, we randomly selected 30 one-cubic-millimeter regions of interest (ROIs) from plaque-bearing areas for each sample and calculated both vascular and plaque densities. Within AD Sample 1 (C = 3), Aβ plaque density and vascular density were significantly positively correlated (r = 0.71465, P < 0.0001). Similar positive correlation was also observed within AD Sample 2 (C = 2; r = 0.60549, P = 0.0004) (Figure 4E). In a descriptive comparison of the basolateral complex of the amygdala, vascular density in AD Sample 1 was approximately threefold higher than that in the control block (Extended Data Figure 6). These analyses identify positive within-specimen spatial associations between Aβ plaque density and vascular density in the two AD specimens examined and demonstrate the use of FIDELITY for millimeter-scale quantification of disease-associated spatial relationships in archived human FFPE tissue.

### FIDELITY supports five rounds of cyclic immunolabeling in the same specimen

Cyclic immunolabeling has been employed to achieve high-plex protein imaging for revealing complex tissue microenvironments ^6,7,44^. To determine whether FIDELITY supports repeated immunolabeling, we adapted the photobleaching-based iterative labeling strategy used in the HIF-Clear+ protocol ^26^. As a proof of concept, we completed five sequential rounds of immunolabeling on a single FFPE mouse brain over a two-month experimental period (Extended Data Figure 7), and did not observe noticeable tissue distortion and swelling (Figure 5A). In each round, the brain was labeled with two antibodies and DiO as a structural reference. Figure 5B shows representative labeling patterns and the subcellular details of the 10 target proteins in both XY and XZ orientations. The 10 datasets were subsequently merged using DiO as the structural reference to visualize their combined spatial distributions (Figure 6A-B, Supplementary video 3).

**Figure 5.**
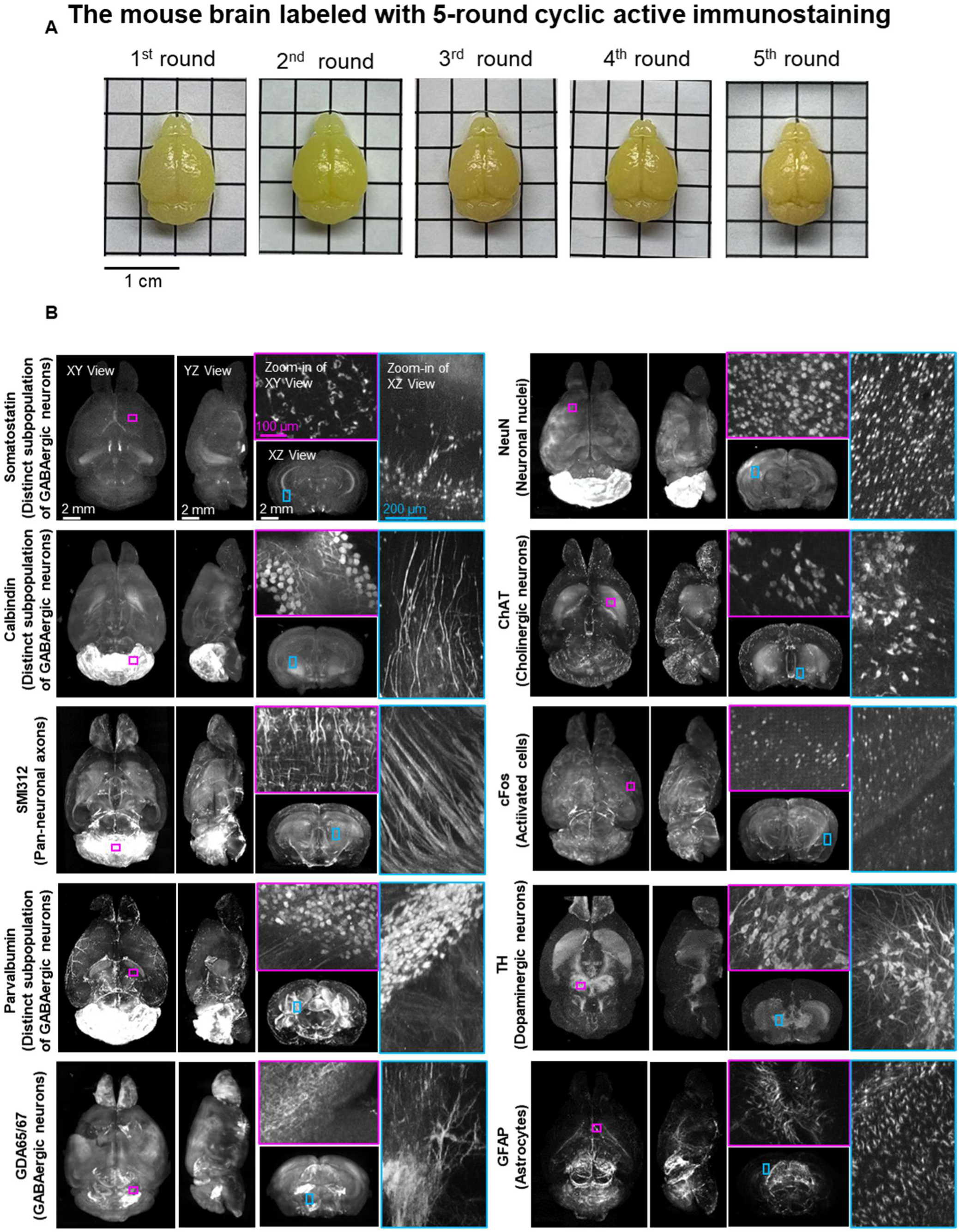
10-plex whole mouse brain imaging enabled with 5-rounds of active cyclic immunolabeling after FIDELITY pipeline. (A) Macroscopic images of the iteratively labeled mouse brain following each round of immunostaining. The same FFPE mouse brain was imaged after each of five rounds of immunostaining. Images show consistent morphology and no visible tissue swelling or distortion across staining cycles. (B) Light-sheet microscopy imaging datasets of three-round immunolabeling performed on one whole FFPE mouse brain. Projection images of the horizontal (XY), sagittal (YZ), and transverse (XZ) views are shown. Projection images (100 μm thick) of magenta-lined and cyan-lined regions marked in the XY and XZ views are magnified and displayed within correspondingly colored frames. Scale bars are indicated in the first row. FFPE, formalin-fixed paraffin-embedded.

**Figure 6.**
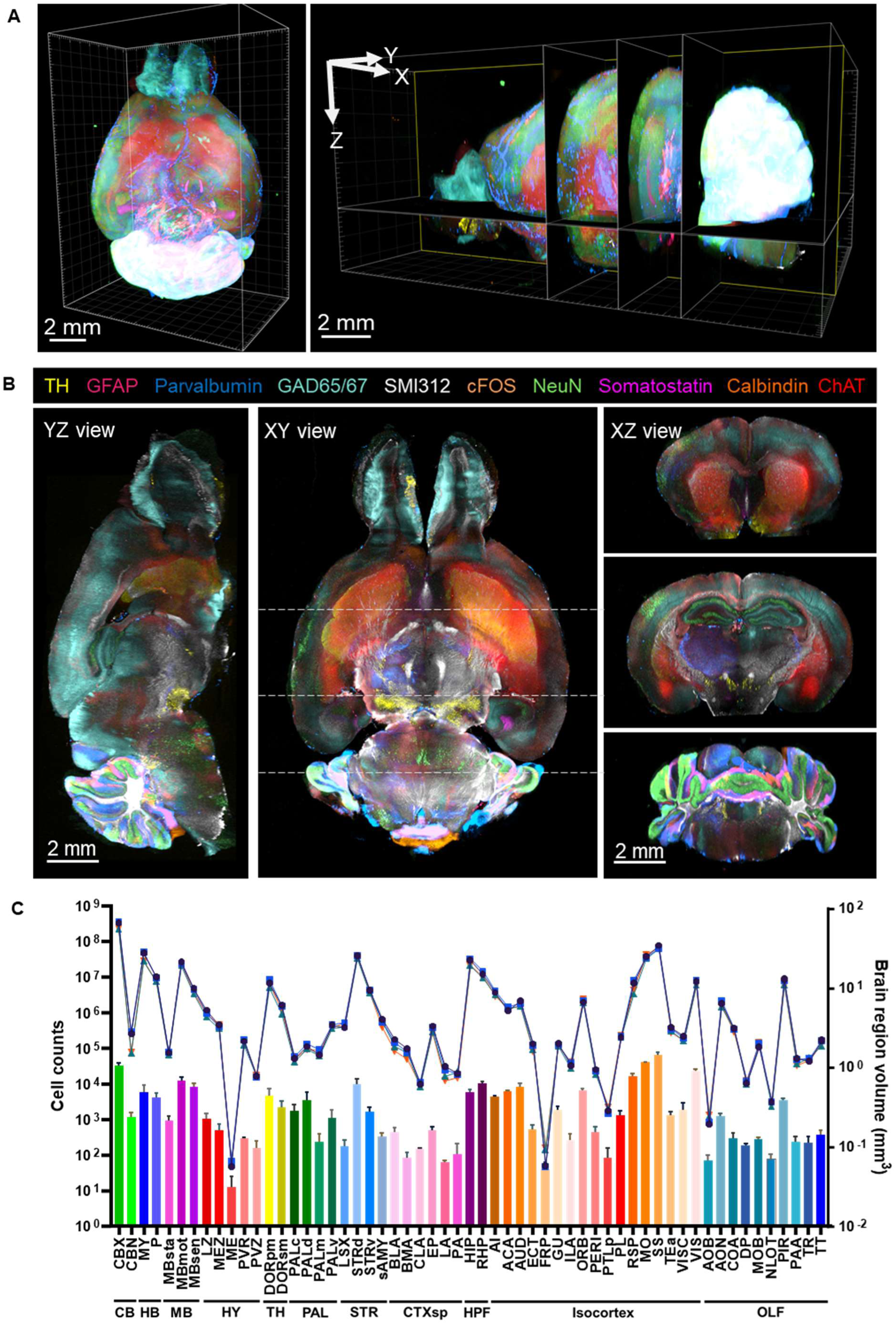
Registration of FIDELITY multi-round staining datasets enables multi-channel 3D visualization and whole-brain cell profiling. (A) (B) Merged multichannel image generated from the datasets in Figure 4. Optical sections are shown in horizontal (XY) and sagittal (YZ) views. The Transverse (XZ) views show optical sections corresponding to the positions indicated by dotted lines in the XY view. (C) Quantification of the volumes of major anatomical brain regions in ABA and the corresponding counts of parvalbumin (PV)-positive neurons (n = 4 samples). Brain region volumes are plotted individually as line graphs, whereas PV cell counts are shown as mean ± SD in the bar graph.

To assess time efficiency, we compared one-time tissue preparation times and cumulative staining times corresponding to five single-round equivalents (Extended Data Figure 8). Single-round staining time included antibody incubation time for a whole mouse brain and associated washing steps. The tissue processing plus cumulative staining time across the five-round FIDELITY experiment totaled 9 days, whereas protocol-derived estimates for INSIHGT, CUBIC-HV, and iDISCO were 21.5, 39, and 77 days, respectively ^20,45,46^. For the comparator methods, these estimates were calculated by multiplying the reported single-round staining durations for whole-mouse-brain specimens by five. These values represent cumulative staining times rather than the total duration of five-round cyclic workflows. Under the specified labeling conditions, FIDELITY required less cumulative staining time than the comparator protocols. Importantly, its compatibility with active electrophoretic staining broadens the choice of labeling strategies for large FFPE specimens, allowing rapid staining protocols to be incorporated into repeated molecular profiling.

### FIDELITY-processed mouse brains support atlas registration and regional neuronal profiling

Because tissue distortion can hinder automated registration of cleared-brain datasets ^47^, we evaluated whether FIDELITY preserved brain morphology sufficiently for standardized anatomical analysis. Four FIDELITY-processed hemispheres were imaged and registered to the Allen Brain Atlas (ABA) using Bi-channel Image Registration and Deep-learning Segmentation (BIRDS).^48^ Major brain-region volume profiles were nearly superimposable across the four datasets, indicating highly consistent morphology across FIDELITY-processed samples (Figure 6C, the line graphs). The estimated volumes were also similar to published measurements obtained from SOLID-cleared brains and conventional serial sections ^14,49^ (Extended Data Figure 9A), providing an external benchmark for atlas-based volumetry.

We next quantified parvalbumin-positive interneurons across registered brain regions (Figure 6C, the bar graph). Regional distribution patterns were similar to those reported previously ^49^, although the absolute cell numbers were lower (Extended Data Figure 9B). Together, these results demonstrate that FIDELITY reproducibly preserves whole-brain morphology across specimens, enabling consistent registration to a standardized atlas and reliable regional quantification of both anatomical volumes and neuronal populations.

### FIDELITY resolves spatially distinct cellular compositions within a human glioma specimen

We further applied FIDELITY to a surgically resected human glioma specimen. H&E staining before FIDELITY revealed spatially distinct dense and loose regions (Extended Data Figure 10A). After clearing (Extended Data Figure 10B), we performed two rounds of immunolabeling targeting five neural and glial markers (Extended Data Figure 10C and D). Three-dimensional visualization showed markedly higher NeuN and GFAP labeling in the dense region, indicating substantial differences in cellular composition between the two compartments.

Co-expression of the oligodendroglial transcription factor Olig2 and the astrocytic marker GFAP has been implicated in tumor lineage plasticity and therapy resistance ^50–52^. Given this association, we quantified the density of Olig2⁺, GFAP⁺, and Olig2⁺/GFAP⁺ double-positive cells (Extended Data Figure 10E). The dense region contained approximately 2.0 × 10⁴ GFAP⁺ cells mm⁻³ and 1,309 Olig2⁺/GFAP⁺ cells mm⁻³, whereas the loose region contained approximately 3.4 × 10² GFAP⁺ cells mm⁻³ and 21 double-positive cells mm⁻³. Thus, the densities of both GFAP⁺ and Olig2⁺/GFAP⁺ cells were approximately 60-fold higher in the dense region. In contrast, Olig2⁺ cell density did not differ significantly between the dense and loose regions (8.7 × 10⁴ cells mm⁻³ and 6.9 × 10⁴ cells mm⁻³, respectively) (Extended Data Figure 10F, left panel).

Despite these differences in absolute cell density, Olig2⁺/GFAP⁺ cells represented approximately 6% ± 0.3% of the GFAP⁺ population in both regions (Extended Data Figure 10F, right panel). These data identify marked compartment-specific differences in GFAP⁺ and Olig2⁺/GFAP⁺ cell density while showing a similar relative abundance of Olig2⁺/GFAP⁺ cells within the GFAP⁺ population. Together, this analysis demonstrates that FIDELITY can resolve spatially distinct cellular compositions and marker co-expression patterns within an intact human glioma specimen.

## Discussion

In this study, we developed FIDELITY, an epoxy-free workflow that balances optical transparency, immunostaining performance, and tissue integrity for large-volume FFPE imaging. The CSR-optimized SDS–glycine formulation integrates delipidation and epitope retrieval while preserving tissue stiffness, replacing the prolonged epoxy-reinforcement step and high-temperature antigen retrieval used in HIF-Clear with a single aqueous pretreatment. Importantly, this physicochemical balance maintains compatibility with active electrophoretic staining, broadening the choice of labeling strategies for large FFPE specimens and enabling rapid antibody delivery in cyclic workflows. These features supported volumetric imaging and quantitative analysis of archived human Alzheimer’s disease tissue while retaining compatibility with conventional histopathology. Meanwhile, five-round immunolabeling in a centimeter-scale FFPE mouse brain showed that repeated molecular imaging could be achieved with minimal morphological distortion, while consistent atlas-based measurements across specimens supported reproducible anatomical analysis. Two-round immunolabeling of a surgically resected human glioma specimen further extended this cyclic capability to human pathological tissue. Together, these findings establish FIDELITY as a practical platform for cyclic molecular profiling of large FFPE specimens within their preserved 3D spatial context.

Beyond archived FFPE human specimens, FIDELITY also provides a practical workflow for experimental animal tissues. In the standard workflow, paraffin embedding allows specimens to be archived before subsequent volumetric analysis. However, long-term storage is not required for every application. Our observation that tissue transparency was maintained when paraffin embedding was omitted suggests that the workflow could be further simplified for freshly collected animal specimens. Further validation of immunostaining performance, tissue stiffness, and compatibility with cyclic labeling is needed to determine whether this modified workflow retains the broader capabilities of the standard FIDELITY protocol.

The design of tissue-clearing workflows must account for the preservation requirements of the intended application. Recently published SeeDB-Live illustrates this principle in living tissues by combining refractive-index matching with isotonic conditions to support structural and functional imaging ^53^. In FIDELITY, the corresponding challenge is to improve molecular accessibility while preserving the mechanical stability required for repeated electrophoretic staining of FFPE specimens. The CSR framework, combined with OACD, modeled immunostaining performance across SDS– glycine combinations as a response surface, enabling the identification of an optimized formulation without relying solely on one-factor-at-a-time titration. However, the present optimization was limited to two chemical factors and primarily focused on immunostaining performance. Future CSR-based studies could incorporate additional chemical and process variables, including pH, temperature, incubation time, and electrophoretic conditions, while jointly evaluating multiple responses, such as immunostaining performance, optical transparency, and Young’s modulus. Such multi-response optimization could support more systematic physicochemical balancing of molecular accessibility, optical transparency, and mechanical stability.

Within the FIDELITY pipeline, we utilized eFLASH ^24^ for its rapid, uniform staining. However, its reliance on specialized instrumentation and restricted sample size may limit broader adoption. Alternative deep-labeling chemistries such as INSHIGT, which leverages chaotropes and macromolecular diffusiophoresis to achieve staining homogeneity with broad antibody compatibility ^46^, may provide a complementary solution, although its compatibility with FIDELITY-treated FFPE specimens requires further testing.

Extending FIDELITY to pathology enables volumetric, quantitative readouts surpassing conventional 2D histology. In Alzheimer’s disease (AD), vascular dysfunction and contradictory increased angiogenesis have been reported, yet vessel density changes have been inconsistent across cohorts and stages^41,54^. Conventional thin sections may undersample tortuous structures such as microvasculature and perivascular spaces resulting in inaccurate quantification. In this context, FIDELITY provides 3D metrics, including plaque volumes, plaque–vessel distance, capillary length density, and branching complexity, reducing sectioning bias and examine structural changes accurately. FIDELITY’s translational potential could extend beyond AD to other diseases, such as spatial mapping of α-synuclein and cholinergic pathology in Parkinson’s disease ^55–57^.

In summary, FIDELITY is a robust and versatile platform for multiplexed 3D pathology of archived FFPE specimens. By interoperating with emerging imaging and molecular-profiling technologies, it has the potential to transform spatial disease research and broaden translational applications.

## Materials and Methods

### Experimental animals and human samples

We used 8-week-old C57/B6 male mice to develop the FIDELITY pipeline. All animal procedures and handling complied with guidelines from the Institutional Animal Care and Use Committee of Academia Sinica (IACUC protocol No.: 24-08-2241), Taiwan. The analyses involving human participants were reviewed and approved by the Institutional Review Board of National Taiwan University Hospital (IRB No.: 202203079RINC), and all participants provided signed informed consent. Archived human FFPE brain blocks from clinically and neuropathologically confirmed Alzheimer’s disease (AD) patients and age-matched controls were obtained from the Iowa Neuropathology Resource Laboratory, University of Iowa. All specimens were de-identified and collected under IRB-approved protocols by the provider. Experimental procedures and the associated guidelines for processing archival FFPE human tissue were approved by the Research Ethics Committee of National Tsing Hua University (REC No.: 11312HM202).

### Sample collection and preparation

For mouse brain samples, the mice were euthanized by an injection of 0.2 mL of a 30% Urethane saline solution. The euthanized mice were perfused with 20 mL of cold phosphate-buffered saline (PBS), followed by 20 mL of 4% PFA. The collected brains were further fixed using 4% PFA at room temperature for 2-7 days before paraffin processing, or at 4°C for 24 hours before SHIELD processing. Human brain tissue specimens were surgically collected from the right temporal lobe of patients with glioma and fixed with 4% PFA at room temperature for 2-7 days. For paraffin embedding, fixed specimens were dehydrated and processed in a tissue processor (Tissue-Tek VIP 5 Jr., Sakura Finetek Japan Co. Ltd., Japan) under vacuum conditions and then embedded in paraffin wax. For the human brain FFPE blocks provided by the Iowa Neuropathology Resource Laboratory, tissues were fixed in 10% neutral buffered formalin for a minimum of 14 days, then sectioned and placed into cassettes. Sections were processed on a Tissue-Tek VIP tissue processor in a series of graded ethanol, xylene, and paraffin wax. Tissues were then embedded using Histoplast PE paraffin wax from Epredia on an Epredia Histostar embedding station.

### Deparaffination and rehydration for whole FFPE blocks

FFPE blocks were incubated at 65-70 °C to melt the paraffin. Residual paraffin was removed through immersion in xylene for 24 hours, with the xylene being changed at least twice. Dewaxed tissues were washed twice in absolute ethanol for 4 hours each to remove residual xylene, followed by sequential rehydration in 95%, 85%, 75%, and 55% ethanol (diluted with distilled water) for at least 3 hours per step. Rehydrated tissues were then rinsed in PBS. When necessary, tissues were stored temporarily in PBSN (PBS containing 0.02% sodium azide) at 4 °C.

### Tissue preparation for atomic force microscopy and transmission spectrum measurements

FFPE mouse brains were dewaxed and rehydrated as described above. Rehydrated brains were sectioned into 1-mm-thick slices using a vibratome (VT1200S, Leica, Germany). Ten brain slices were incubated separately in ten test solutions at 54 °C for 16 hours. The test solutions were PBS, and 10% (v/v) PBS solutions of Triton X-100, Tween 20, sodium cholate, NP-40, DMSO, CHAPS, SDS, urea, and glycine. After incubation, the slices were thoroughly washed with PBS.

### Atomic force microscopy (AFM)

Samples were prepared as described above. Young’s modulus was measured using a bio-AFM system (NanoWizard® 3, JPK, Bruker Corp., Billerica, MA, USA). A spherical borosilicate glass probe (CP-PNP-TR, radius = 10 µm) with a reflective Cr/Au coating was used for indentation experiments. For elastic modulus mapping (Figures 1A and 1F), samples were scanned in Quantitative Imaging (QI™) mode under the Advanced Imaging setting. Each scan covered a 10 µm × 10 µm area with a resolution of 64 × 64 pixels. The vertical travel range (z-length) was set to 2 µm, the maximum applied force (setpoint) to 1.2 nN, and the dwell time per force curve to 40 ms.

For regional elasticity measurements (Figure 2C), experiments were performed in contact mode using force spectroscopy. Within the striatal region of FFPE mouse brain sections, ten randomly selected areas were probed. In each area, a 2 × 2 grid with 10 µm spacing was defined, and force–distance curves were recorded at each grid point. Indentation parameters were as below: maximum applied force = 1.2 nN, indentation depth = 2 µm, and loading speed = 2 µm s⁻¹. After each force curve, the cantilever was fully retracted to ensure complete disengagement from the sample surface, minimizing residual adhesive or lateral forces.

Data were analyzed using the manufacturer’s software (JPK Data Processing, version 6.1.203, Bruker).

Force–distance curves were fitted to the Hertzian contact model to extract the apparent local Young’s modulus. The tissue was modeled as an incompressible material, with the Poisson’s ratio fixed at 0.5 for all analyses.

### Transmission Spectrum Measurement

Samples were prepared as described above. The transmission spectrum was measured using a broadband white-light LED as the illumination source and a high-resolution triple-grating spectrometer (iHR550, Horiba). The LED provided stable output across the visible range, suitable for broadband spectral characterization. The emitted light was slightly focused through an optical system to reduce the beam diameter while maintaining uniform intensity across the sample area, thereby avoiding angular dispersion or illumination artifacts.

Transmitted light was collected and directed into the spectrometer entrance slit (spectral resolution = 0.025 nm). The optical configuration was optimized to suppress stray light and maintain a high signal-to-noise ratio. Samples were mounted between two microscope slides to ensure consistent optical alignment and minimize reflection as well as scattering.

To obtain normalized transmittance spectra, a two-step measurement was performed. A reference spectrum was first recorded with blank slides only, followed by measurement of the sample spectrum under identical conditions. Transmittance was calculated as the ratio of sample to reference spectra, compensating for the spectral response of the source and optical components and ensuring reproducible, high-accuracy results.

### Immunofluorescent staining of FFPE sections

FFPE mouse brain sections (5 µm thick) were cut and mounted onto MAS-GP hydrophilic adhesion slides (S9901, Matsunami Glass Ind., Ltd., Osaka, Japan). Sections were dewaxed and rehydrated, then incubated in the test solutions of interest at 54 °C for 16 hours. After being washed thoroughly with PBS, sections were blocked in PBS containing 0.2% BSA and 0.2% normal donkey serum for 30 minutes at room temperature.

Primary antibodies including SMI312 (837904, BioLegend, CA, USA; 1:500 dilution) and NeuN (24307S, Cell Signaling Technology, MA, USA; 1:200 dilution) were diluted in blocking buffer and applied at 37 °C for 1 hour. After being washed with PBS, sections were incubated with secondary antibodies containing donkey anti-rabbit Alexa 488 Fab (711-547-003, Jackson ImmunoResearch, ME, USA; 1:500 dilution) and donkey anti-mouse Rhodamine X red Fab (715-297-003, Jackson ImmunoResearch, ME, USA; 1:500 dilution) for 1 hour at room temperature.

Sections were washed again with PBS, counterstained with DAPI (5 µM in PBS), and mounted in 50% glycerol. Imaging was performed using a multipoint confocal microscope (Dragonfly 200, Andor, Oxford Instruments, UK) equipped with a 20× 0.75 NA UPlanSApo objective (Evident Corp., Tokyo, Japan).

### Complex system response (CSR)-guided optimization of delipidation parameters

We employed a customized MATLAB application to perform CSR fitting and multi-objective optimization ^37^. The parameter ranges were defined as 4–10% for SDS concentration and 2–10% for glycine concentration. An Orthogonal Array Composite Design (OACD) table was generated to prepare calibration solutions for delipidation optimization (Supplementary Table 1). Each test solution was applied to FFPE sections prior to immunostaining as described above. Fluorescence signal-to-background (S/B) ratios were quantified using Fiji software ^58^. For each staining condition, 120 individual cells were randomly selected across multiple regions of interest. The mean fluorescence intensity of each cell was measured using circular regions of interest (ROIs) drawn around the labeled cell bodies. Background intensity was determined by measuring the mean fluorescence within an adjacent cell-free area of identical size. The S/B ratio for each cell was calculated as the ratio of mean signal intensity to mean background intensity. All measurements were performed on raw, unprocessed images with identical acquisition parameters. The extrema were identified by importing the OACD table along with the corresponding immunolabeling outcomes into the MATLAB framework.

### SHIELD processing, SDS-electrophoretic delipidation

PFA-fixed specimens were incubated in SHIELD-OFF solution at 4 °C for 96 hours, followed by incubation for 24 hours in SHIELD-ON solution at 37 °C. All reagents were prepared using SHIELD kits (LifeCanvas Technologies, Seoul, South Korea) according to the manufacturer’s instructions. For SDS-electrophoretic delipidation, SHIELD-processed specimens were placed in a stochastic electro-transport machine (SmartClear Pro II, LifeCanvas Technologies, Seoul, South Korea) running at a constant current of 1.2 A for 5 days.

### FLASH delipidation

The FFPE mouse brain blocks were dewaxed and rehydrated as described above, followed by incubation in FLASH reagent (4% w/v SDS, 200 mM borate) at 54 ℃ for 18 hours.^30^. The delipidated specimens were washed with PBST (1% Triton X-100 in PBS) at room temperature for at least 1 day.

### FIDELITY pipeline

The FIDELITY pipeline is illustrated in Figure 3 and described in detail below.

#### Deparaffination and rehydration

FFPE blocks were dewaxed and rehydrated as described above.

#### Decolorization (optional)

Non-perfused or pigment-rich specimens were decolorized to minimize light absorption during imaging. Tissues were incubated in 10% H₂O₂ (in PBS) at 4 °C for 16–72 hours, depending on tissue size and pigment deposition. The H₂O₂ solution was changed every 24 hours.

#### Optimized delipidation/antigen retrieval

Specimens were incubated in FIDELITY solution (6.9% SDS and 5.5% Glycine in 200 mM Boric acid, pH 7.0) for 16 hours at 54 °C. The specimens were then washed thoroughly with PBST for at least 24 hours at 37 °C with the PBST being changed twice.

#### Electrophoretic immunolabeling (active staining)

The procedure was modified from the previously published eFLASH protocol ^24^ and was conducted in a SmartLabel System (LifeCanvas Technologies, Seoul, South Korea). The specimens were preincubated overnight at room temperature in sample buffer (240 mM Tris, 160 mM CAPS, 20% w/v D-sorbitol, 0.9% w/v sodium deoxycholate). Each preincubated specimen was placed in a sample cup (provided by the manufacturer with the SmartLabel System) containing primary, corresponding secondary antibodies and DiO diluted in 8 mL of sample buffer. Information on antibodies, DiO and their optimized quantities is detailed in Supplementary Table 2. The specimens in the sample cup and 500 mL of labeling buffer (240 mM Tris, 160 mM CAPS, 20% w/v D-sorbitol, 0.2% w/v sodium deoxycholate) were loaded into the SmartLabel System. The device was operated at a constant voltage of 90 V with a current limit of 400 mA. After 18 hours of electrophoresis, 300 mL of booster solution (20% w/v D-sorbitol, 60 mM boric acid) was added, and electrophoresis continued for 4 hours. During the labeling, the temperature inside the device was kept at 25 ℃. Labeled specimens were washed twice (3 hours per wash) with PTwH (1× PBS with 0.2% w/v Tween-20 and 10 μg/mL heparin),^[23]^ and then post-fixed with 4% PFA at room temperature for 1 day. Post-fixed specimens were washed twice (3 hours per wash) with PBST to remove any residual PFA.

#### Refractive index (RI) matching

Before imaging, specimens were immersed sequentially in 50% NebClear and pure NebClear (RI = 1.519; Nebulum, Taipei, Taiwan) for one day each at room temperature to achieve RI matching. Alternatively, RI matching was performed by immersing specimens in a 1:1 dilution of CUBIC-R [28] for one day, followed by pure CUBIC-R for another day.

#### Volumetric imaging and 3D visualization

For the mouse brains and the human glioma specimen, images were acquired using a light-sheet microscope (SmartSPIM, LifeCanvas Technologies, Seoul, South Korea) with a 3.6x customized immersion objective (NA = 0.2, working distance = 1.2 cm). For the AD human amygdala specimens, imaging was performed using a multipoint confocal microscope (Andor Dragonfly 200, Oxford Instruments, UK) with UMPLFLN10XW objective (10x, NA = 0.3, working distance = 3.5 mm). 3D visualization was performed using Imaris software (Imaris 9.5.1, Bitplane, Belfast, UK).

#### Photobleaching

Immunolabeled and imaged specimens were placed in a multi-well plate and immersed in RI-matching solutions to retain their transparency. The plate was then sealed with paraffin. A 100-W projection lamp with an LED array was placed on the plate to quench fluorescence signals ^26^. In our experience, approximately 18 hours photobleaching is sufficient for a 2-mm-thick sample, whereas 3 days is required for a whole mouse brain (approximately 8 mm to 1 cm in thickness).

### Whole brain parvalbumin (PV) interneuron profiling

For the parvalbumin-positive cell quantification, we used a trained nnU-net model to perform prediction and cell segmentation ^59^. The segmented binary images were then processed with a 3D spatial filter in cellfinder ^60^ to obtain the coordinates of the cell centers. To assign the detected cells to the corresponding brain regions, Bi-channel Image Registration and Deep-learning Segmentation (BIRDS ^48^) was used to register the CCFv3 atlas provided by the Allen Brain Institute, to the autofluorescence channel, which was adapted by cellfinder.

### Multichannel image registration

In multiplex immunolabeling of whole mouse brains, each dataset contained one structural reference channel labeled with DiO and one antibody-stained channel. The first-round dataset served as the standard brain. Image registration was performed using the Advanced Normalization Tools (ANTs) package ^61^. All datasets were downsampled by a factor of 5 in the XY plane and 10 in the Z axis and preprocessed by denoising, N4 bias-field correction, histogram matching, and intensity truncation. Spatial transforms were estimated using the antsRegistrationSyN[s] workflow, with linear (iterations 1000–100) and nonlinear SyN stages (CC metric, step size 0.15). Transformation parameters derived from DiO reference channels were applied to the antibody-stained channels. Registered datasets were merged and visualized using Imaris (Bitplane, Zurich, Switzerland).

### Conventional histology and imaging

To assess tissue morphology and evaluate the compatibility of FIDELITY-processed samples with conventional histopathology, hematoxylin and eosin (H&E) staining was performed on FFPE sections. FFPE tissue sections (5 µm thick) were prepared using a rotary microtome (BIO CUT, Leica Biosystems, Nussloch, Germany) and mounted onto MAS-GP hydrophilic adhesion slides (S9901, Matsunami Glass Ind., Ltd., Osaka, Japan). H&E staining was carried out using automated workstations (ST5010 XL and CV5030, Leica Biosystems, Nussloch, Germany). Whole-slide images were acquired with a 3DHISTECH Pannoramic 250 digital slide scanner (3DHISTECH Kft., Budapest, Hungary) equipped with a 40× / 0.95 NA Plan-Apochromat objective (Zeiss, Oberkochen, Germany).

### Aβ burden and vascular density quantification in AD specimens

To assess the relationship between Aβ plaques and vascular density, 30 regions of interest (ROIs; 1 mm³ each) were randomly selected within plaque-rich areas of each specimen. Both plaque and vascular structures were segmented using the Surface module in Imaris software (version 9.5.1, Bitplane, Belfast, UK) with automatic thresholding based on fluorescence intensity and a minimum object size of 50 µm³ to exclude noises. Surface smoothing was set to 3 µm to preserve structural continuity while minimizing edge artifacts. Aβ plaque and vascular volume densities were then quantified from the segmented surfaces. Pearson’s correlation coefficients were calculated in GraphPad Prism 9.5.1 (GraphPad Software, San Diego, CA, USA).

### Statistics

All statistical analyses and graphing were performed using GraphPad Prism version 9.5.1 (GraphPad Software, San Diego, CA, USA). Data are presented as mean ± standard deviation (SD) unless otherwise indicated. Statistical significance was determined by one-way ANOVA followed by Tukey’s multiple comparison test, with α = 0.05 (*P < 0.05; **P < 0.01; ***P < 0.001; ****P < 0.0001).

## Acknowledgements

We thank the Brain Research Center, National Tsing Hua University, Taiwan; the Pathology Core at the Institute of Biomedical Sciences, Academia Sinica, Taiwan; and the Research Center of Applied Sciences, Academia Sinica, Taiwan, for their technical assistance. We thank Mr. Zhe-Wei Xu for his assistance with parvalbumin-positive cell quantification. We are also grateful to Prof. Chih-Ming Ho and Dr. Wen-Jun Chen for their kind support in CSR analysis. This work was supported by the National Science and Technology Council (NSTC-113-2311-B-007-013, NSTC-114-2314-B-002-299, and NSTC-114-2321-B-A49-014) and by the Brain Research Center under the Higher Education Sprout Project, co-funded by the Ministry of Education and the National Science and Technology Council in Taiwan.

## Data Availability Statement

All image data supporting the findings of this study are included in the main figures and Supplementary Information. Source data for Figures 1D, 1G, 2D, 2E, 4E, 6C, and Extended Data Figures 6C, 9, and 10F are provided in Supplementary Table 3. The original raw image datasets are available for research purposes upon reasonable request from Dr. Li-An Chu

## References

1. Donczo, B. & Guttman, A. Biomedical analysis of formalin-fixed, paraffin-embedded tissue samples: The Holy Grail for molecular diagnostics. J. Pharm. Biomed. Anal. 155, 125–134 (2018).

2. Kokkat, T. J., Patel, M. S., McGarvey, D., Livolsi, V. A. & Baloch, Z. W. Archived Formalin-Fixed Paraffin-Embedded (FFPE) Blocks: A Valuable Underexploited Resource for Extraction of DNA, RNA, and Protein. Biopreserv. Biobank. 11, 101 (2013).

3. Bass, B. P., Engel, K. B., Greytak, S. R. & Moore, H. M. A review of preanalytical factors affecting molecular, protein, and morphological analysis of formalin-fixed, paraffin-embedded (FFPE) tissue: how well do you know your FFPE specimen? Arch. Pathol. Lab. Med. 138, 1520–1530 (2014).

4. Radtke, A. J. et al. IBEX: an iterative immunolabeling and chemical bleaching method for high-content imaging of diverse tissues. Nat. Protoc. 17, 378–401 (2022).

5. Saka, S. K. et al. Immuno-SABER enables highly multiplexed and amplified protein imaging in tissues. Nat. Biotechnol. 37, 1080–1090 (2019).

6. Klevanski, M. et al. Automated highly multiplexed super-resolution imaging of protein nano-architecture in cells and tissues. Nature Communications 2020 11:1 11, 1–11 (2020).

7. Yapp, C. et al. Highly multiplexed 3D profiling of cell states and immune niches in human tumors. Nature Methods 2025 22:10 22, 2180–2193 (2025).

8. Ueda, H. R. et al. Tissue clearing and its applications in neuroscience. Nature Reviews Neuroscience vol. 21 61–79 Preprint at 10.1038/s41583-019-0250-1 (2020).

9. Molbay, M., Kolabas, Z. I., Todorov, M. I., Ohn, T.-L. & ErtUrk, A. A guidebook for DISCO tissue clearing. Mol. Syst. Biol. 17, e9807 (2021).

10. Richardson, D. S. et al. Tissue Clearing. Nature reviews. Methods primers 1, (2021).

11. Tian, T., Yang, Z. & Li, X. Tissue clearing technique: Recent progress and biomedical applications. J. Anat. 238, 489–507 (2021).

12. Yu, T., Zhu, J., Li, D. & Zhu, D. Physical and chemical mechanisms of tissue optical clearing. iScience 24, 102178 (2021).

13. Ke, M. T., Fujimoto, S. & Imai, T. SeeDB: a simple and morphology-preserving optical clearing agent for neuronal circuit reconstruction. Nature Neuroscience 2013 16:8 16, 1154–1161 (2013).

14. Zhu, J. et al. SOLID: minimizing tissue distortion for brain-wide profiling of diverse architectures. Nature Communications 2024 15:1 15, 1–17 (2024).

15. Chung, K. et al. Structural and molecular interrogation of intact biological systems. Nature 497, 332–337 (2013).

16. Tainaka, K. et al. Chemical Landscape for Tissue Clearing Based on Hydrophilic Reagents. Cell Rep. 24, 2196–2210.e9 (2018).

17. Susaki, E. A. et al. Whole-brain imaging with single-cell resolution using chemical cocktails and computational analysis. Cell 157, 726–739 (2014).

18. Chen, Y. et al. Three-dimensional imaging and quantitative analysis in CLARITY processed breast cancer tissues. Sci. Rep. 9, 1–13 (2019).

19. Nojima, S. et al. CUBIC pathology: Three-dimensional imaging for pathological diagnosis. Sci. Rep. 7, 1–14 (2017).

20. Renier, N. et al. IDISCO: A simple, rapid method to immunolabel large tissue samples for volume imaging. Cell 159, 896–910 (2014).

21. Tanaka, N. et al. Mapping of the three-dimensional lymphatic microvasculature in bladder tumours using light-sheet microscopy. British Journal of Cancer 2018 118:7 118, 995–999 (2018).

22. Tanaka, N. et al. Whole-tissue biopsy phenotyping of three-dimensional tumours reveals patterns of cancer heterogeneity. Nat. Biomed. Eng. 1, 796–806 (2017).

23. Reuss, A. M. et al. aDISCO: A clearing method to enable 3D microscopy of large archival paraffin-embedded human tissue blocks. Sci. Adv. 12, 9926 (2026).

24. Yun, D. H. et al. Uniform volumetric single-cell processing for organ-scale molecular phenotyping. Nat. Biotechnol. 1–12 (2025) doi:10.1038/S41587-024-02533-4; SUBJMETA.

25. Park, Y.-G. et al. Protection of tissue physicochemical properties using polyfunctional crosslinkers. Nat. Biotechnol. 37, 73–83 (2019).

26. Lin, Y.-H. et al. Revealing intact neuronal circuitry in centimeter-sized formalin-fixed paraffin-embedded brain. Elife 13, (2024).

27. Sompuram, S. R., Vani, K., Messana, E. & Bogen, S. A. A Molecular Mechanism of Formalin Fixation and Antigen Retrieval. Am. J. Clin. Pathol. 121, 190–199 (2004).

28. Shi, S. R., Key, M. E. & Kalra, K. L. Antigen retrieval in formalin-fixed, paraffin-embedded tissues: an enhancement method for immunohistochemical staining based on microwave oven heating of tissue sections. J. Histochem. Cytochem. 39, 741–748 (1991).

29. Krenacs, L., Krenacs, T., Stelkovics, E. & Raffeld, M. Heat-Induced Antigen Retrieval for Immunohistochemical Reactions in Routinely Processed Paraffin Sections. Methods Mol. Biol. 588, 103 (2010).

30. Messal, H. A. et al. Antigen retrieval and clearing for whole-organ immunofluorescence by FLASH. Nat. Protoc. 16, 239–262 (2021).

31. Na, M. et al. Sodium Cholate-Based Active Delipidation for Rapid and Efficient Clearing and Immunostaining of Deep Biological Samples. Small Methods 6, (2022).

32. Zhao, S. et al. Cellular and Molecular Probing of Intact Human Organs. Cell 180, 796–812.e19 (2020).

33. Rosas-Arellano, A. et al. A simple solution for antibody signal enhancement in immunofluorescence and triple immunogold assays. Histochem. Cell Biol. 146, 421–430 (2016).

34. Parfitt, G. J. Immunofluorescence Tomography: High-resolution 3-D reconstruction by serial-sectioning of methacrylate embedded tissues and alignment of 2-D immunofluorescence images. Sci. Rep. 9, 1–9 (2019).

35. Pirici, D. et al. Antibody elution method for multiple immunohistochemistry on primary antibodies raised in the same species and of the same subtype. Journal of Histochemistry and Cytochemistry 57, 567–575 (2009).

36. Babu, P. K. V. & Radmacher, M. Mechanics of brain tissues studied by atomic force microscopy: A perspective. Front. Neurosci. 13, 453881 (2019).

37. Chen, W.-J., Wang, C.-C., Lin, S.-C., Yao, D.-J. & Ho, C.-M. Global Optimization of Thin-Film Properties in PECVD System Harnessed by Complex-System-Response Platform. 2025 23rd International Conference on Solid-State Sensors, Actuators and Microsystems (Transducers) 1488–1491 (2025) doi:10.1109/TRANSDUCERS61432.2025.11110793.

38. Yan, L., Khong, J. H., Kostadinov, A., Fuh, J. Y. H. & Ho, C.-M. A ubiquitous transfer function links interacting elements to emerging property of complex systems. https://arxiv.org/abs/2408.03347v5 (2024).

39. Xu, H., Jaynes, J. & Ding, X. Combining two-level and three-level orthogonal arrays for factor screening and response surface exploration. Stat. Sin. 24, 269–289 (2014).

40. Montine, T. J. et al. National Institute on Aging-Alzheimer’s Association guidelines for the neuropathologic assessment of Alzheimer’s disease: a practical approach. Acta Neuropathol. 123, 1–11 (2012).

41. Alvarez-Vergara, M. I. et al. Non-productive angiogenesis disassembles Aß plaque-associated blood vessels. Nat. Commun. 12, 1–16 (2021).

42. Kisler, K., Nelson, A. R., Montagne, A. & Zlokovic, B. V. Cerebral blood flow regulation and neurovascular dysfunction in Alzheimer disease. Nat. Rev. Neurosci. 18, 419–434 (2017).

43. Sweeney, M. D., Sagare, A. P. & Zlokovic, B. V. Blood-brain barrier breakdown in Alzheimer disease and other neurodegenerative disorders. Nat. Rev. Neurol. 14, 133–150 (2018).

44. Ehrenberg, A. J. et al. A manual multiplex immunofluorescence method for investigating neurodegenerative diseases. J. Neurosci. Methods 339, (2020).

45. Susaki, E. A. et al. Versatile whole-organ/body staining and imaging based on electrolyte-gel properties of biological tissues. Nature Communications 2020 11:1 11, 1–22 (2020).

46. Yau, C. N. et al. INSIHGT: an accessible multi-scale, multi-modal 3D spatial biology platform. Nature Communications 2024 15:1 15, 1–20 (2024).

47. Susaki, E. A. & Ueda, H. R. Whole-body and Whole-Organ Clearing and Imaging Techniques with Single-Cell Resolution: Toward Organism-Level Systems Biology in Mammals. Cell Chem. Biol. 23, 137–157 (2016).

48. Wang, X. et al. Bi-channel image registration and deep-learning segmentation (Birds) for efficient, versatile 3d mapping of mouse brain. Elife 10, 1–20 (2021).

49. Bjerke, I. E. et al. Densities and numbers of calbindin and parvalbumin positive neurons across the rat and mouse brain. iScience 24, 101906 (2021).

50. Fukuda, S., Kondo, T., Takebayashi, H. & Taga, T. Negative regulatory effect of an oligodendrocytic bHLH factor OLIG2 on the astrocytic differentiation pathway. Cell Death Differ. 11, 196–202 (2004).

51. Tatsumi, K. et al. Olig2-Lineage astrocytes: A distinct subtype of astrocytes that differs from GFAP astrocytes. Front. Neuroanat. 12, 312356 (2018).

52. Myers, B. L. et al. Transcription factors ASCL1 and OLIG2 drive glioblastoma initiation and co-regulate tumor cell types and migration. Nature Communications 15, 1–21 (2024).

53. Inagaki, S. et al. Isotonic and minimally invasive optical clearing media for live cell imaging ex vivo and in vivo. Nature Methods 2026 23:4 23, 839–853 (2026).

54. Govindpani, K. et al. Vascular Dysfunction in Alzheimer’s Disease: A Prelude to the Pathological Process or a Consequence of It? Journal of Clinical Medicine 2019, Vol. 8, Page 651 8, 651 (2019).

55. Ignatavicius, A., Matar, E. & Lewis, S. J. G. Visual hallucinations in Parkinson’s disease: spotlight on central cholinergic dysfunction. Brain 148, 376–393 (2025).

56. Hart de Ruyter, F. J., et al. α-Synuclein pathology in post-mortem retina and optic nerve is specific for α-synucleinopathies. NPJ Parkinsons Dis. 9, 1–9 (2023).

57. Russo, M. et al. The pharmacology of visual hallucinations in synucleinopathies. Front. Pharmacol. 10, 479989 (2019).

58. Schindelin, J. et al. Fiji: an open-source platform for biological-image analysis. Nature Methods 2012 9:7 9, 676–682 (2012).

59. Isensee, F., Jaeger, P. F., Kohl, S. A. A., Petersen, J. & Maier-Hein, K. H. nnU-Net: a self-configuring method for deep learning-based biomedical image segmentation. Nat. Methods 18, 203–211 (2021).

60. Tyson, A. L. et al. A deep learning algorithm for 3D cell detection in whole mouse brain image datasets. PLoS Comput. Biol. 17, e1009074 (2021).

61. Avants, B. B. et al. A reproducible evaluation of ANTs similarity metric performance in brain image registration. Neuroimage 54, 2033–2044 (2011).

