## Extended data figures for "Physicochemical balancing facilitates fast cyclic immunolabeling in centimeter-scale FFPE tissues"

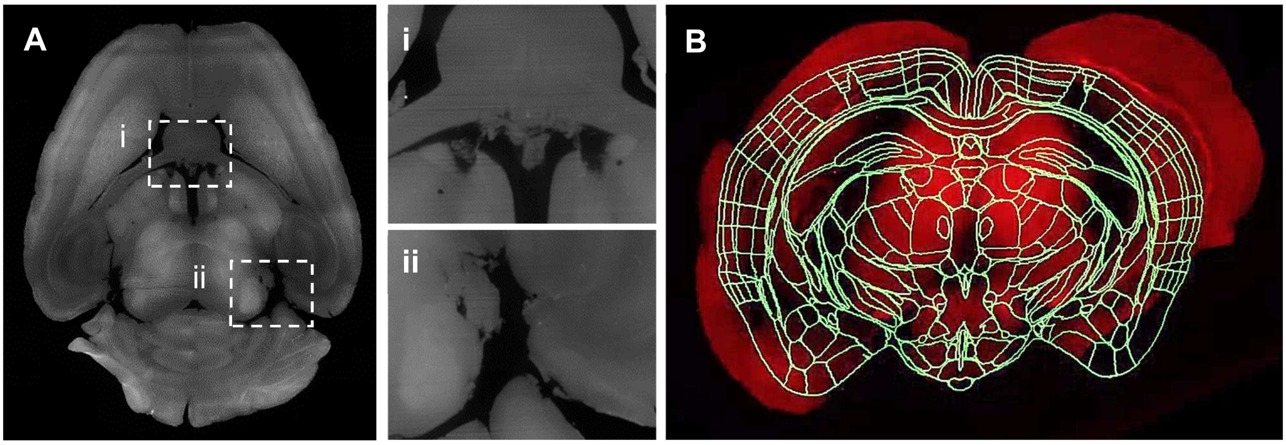


**Extended Data Figure 1. Tissue deformation in epoxy-processed samples during clearing and immunolabeling.**

Degraded epoxy was insufficient to protect brain tissue during harsh electrophoresis, resulting in tissue damage (panels A, A-i and A-ii) and distortion (panel B).

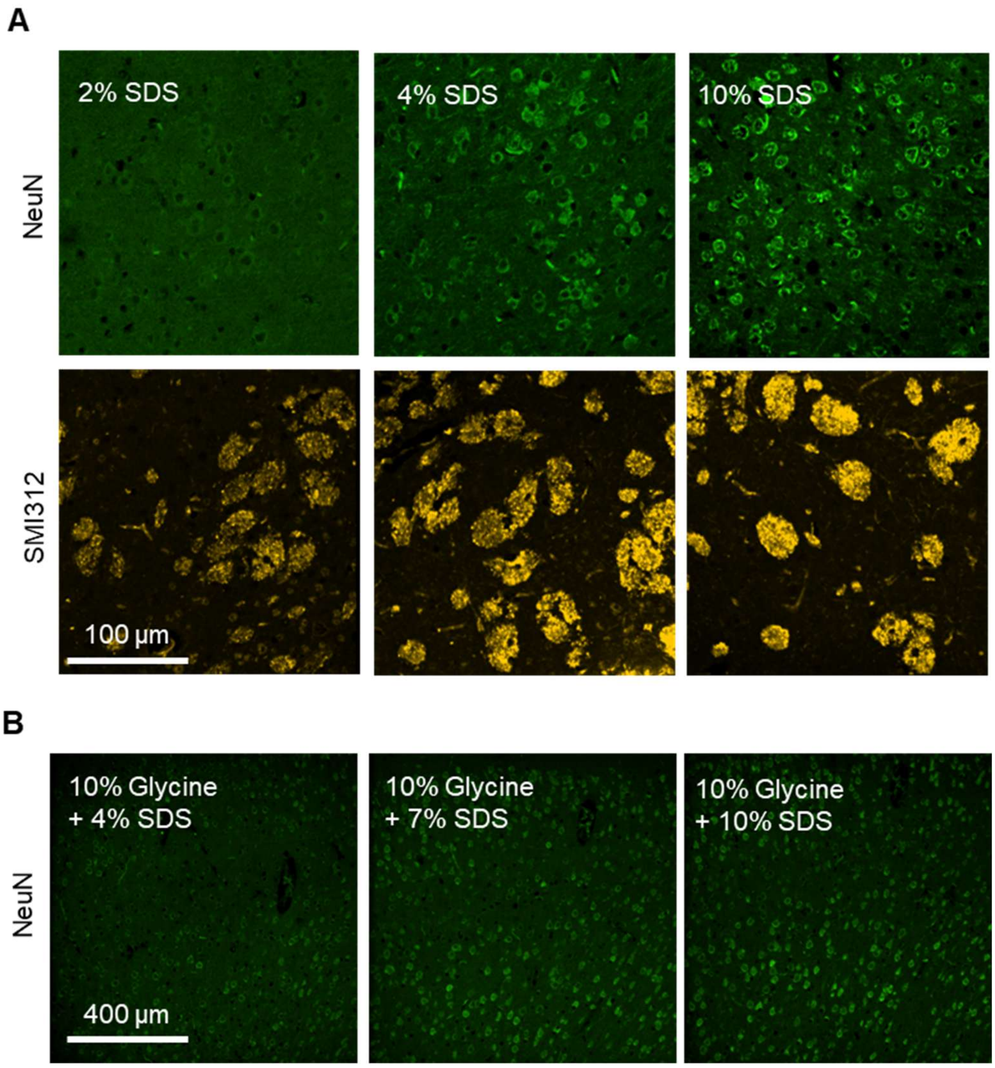


**Extended Data Figure 2. Staining tests of varying glycine and SDS concentrations.**

(A) Dose-dependent effect of SDS on antigen retrieval performance.

(B) Addition of 10% Glycine attenuates the antigen retrieval performance of SDS.

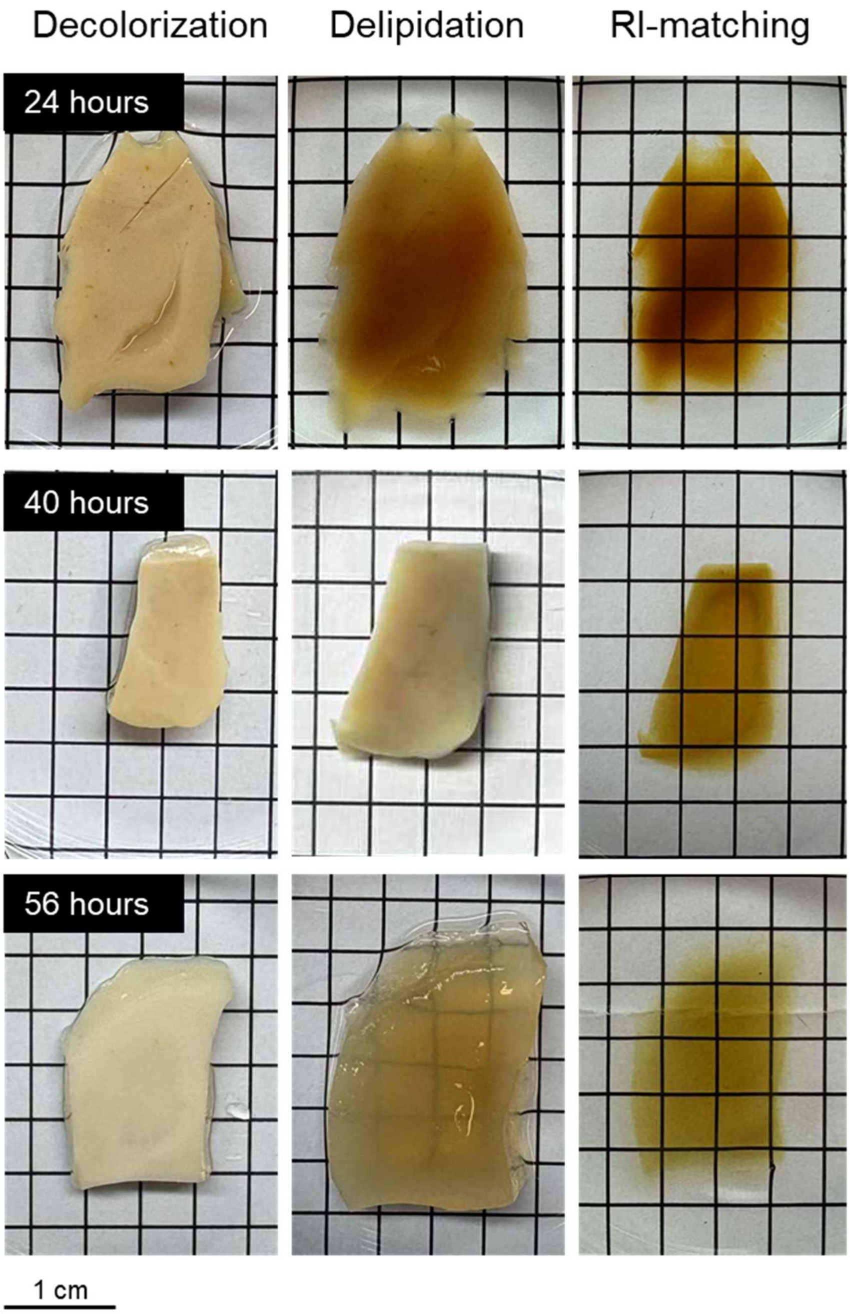


**Extended Data Figure 3. Time-course test of decolorization.** Human brain tissues were retrieved from FFPE blocks by dewaxing and rehydration, and then incubated in 10% $H_2O_2$ at 4 °C for 24, 40, or 56 hours.

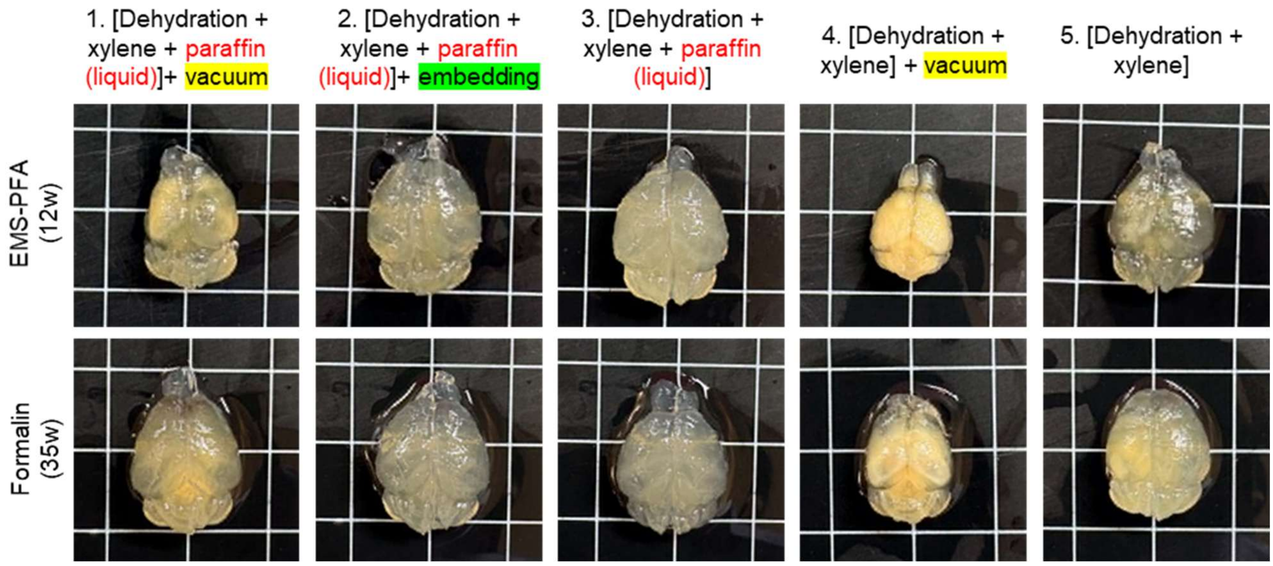


**Extended Data Figure 4. Paraffin embedding is dispensable for FIDELITY processing of freshly collected animal tissues.** Mouse brains fixed with 4% PFA (top) or formalin (bottom) were processed using FIDELITY protocols with or without liquid-paraffin immersion and paraffin embedding. Comparable tissue transparency was achieved under the third condition (the third column), indicating that liquid-paraffin immersion is essential, whereas the subsequent paraffin-embedding step can be omitted for freshly collected specimens when long-term storage is not required.

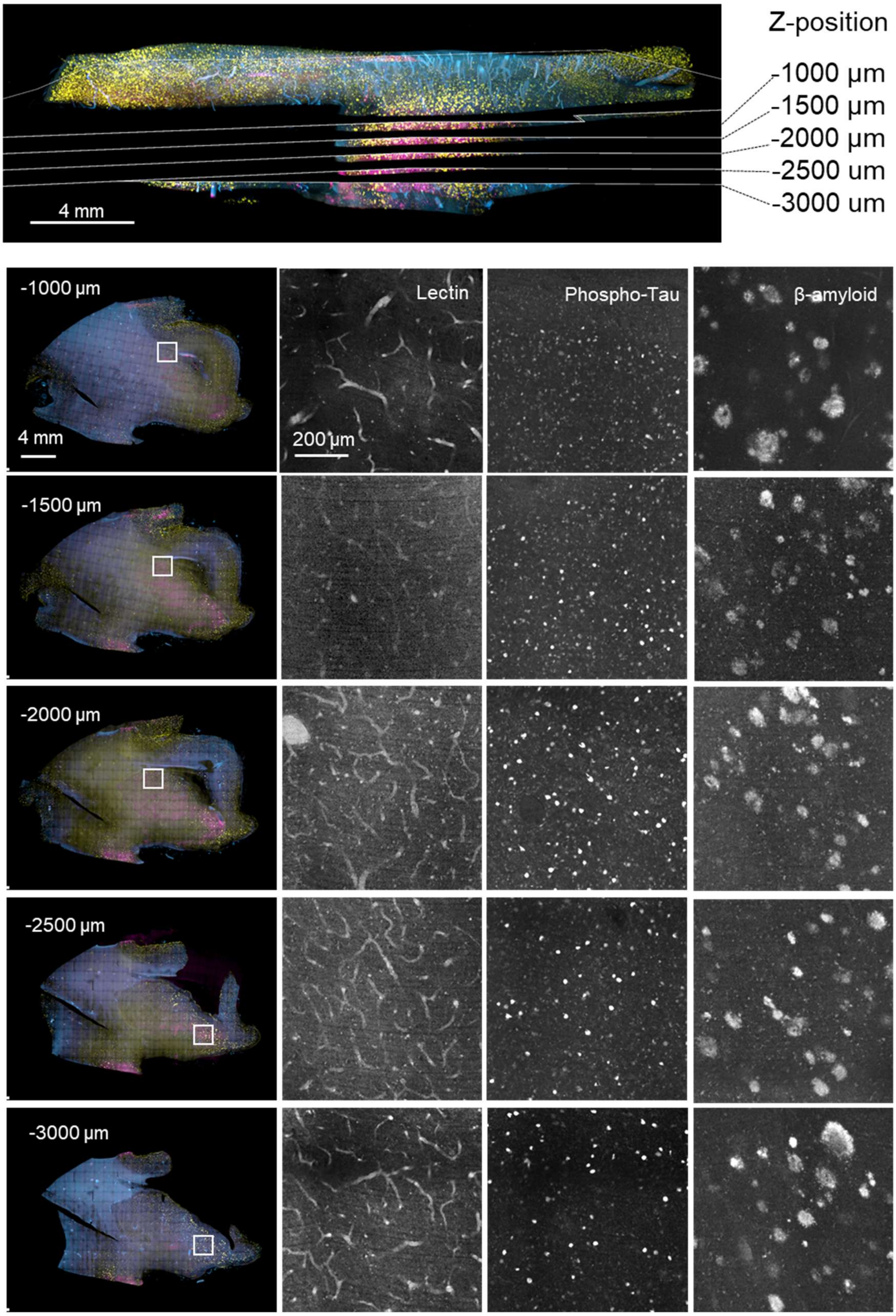


**Extended Data Figure 5. Optical sections at various depths of the cleared FFPE amygdala tissue block from an AD patient shown in Figure 6B.** An overview image indicates the z-positions at which optical sections were acquired. For each z-position, merged fluorescence images are shown, along with single-channel magnified views for lectin (vasculature), phospho-Tau, and β-amyloid**.**

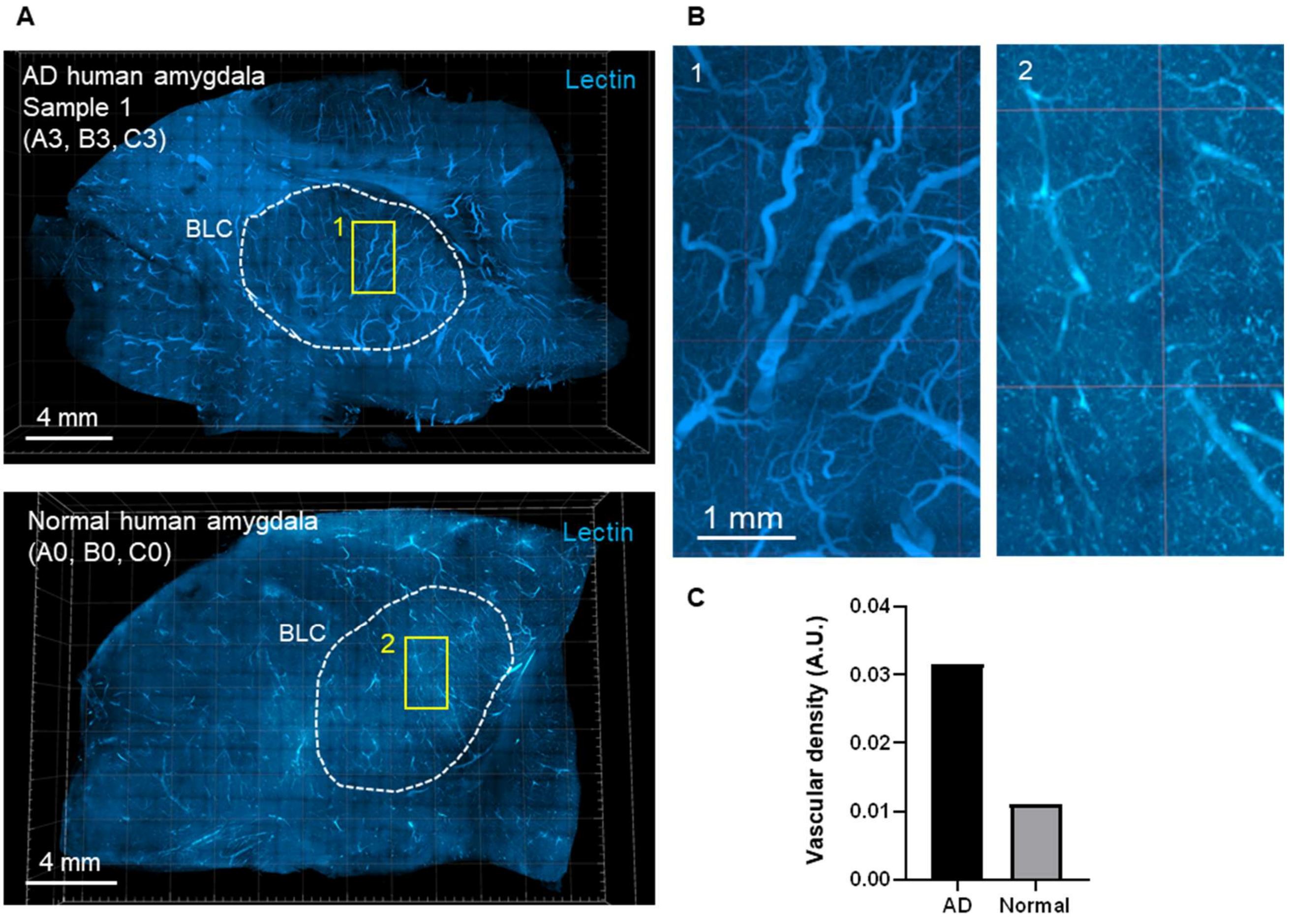


**Extended Data Figure 6. Basolateral amygdala vascular density in AD and normal specimens.**

(A) 3D renderings of FFPE human amygdala specimens, with the basolateral complex (BLC) marked by white-dotted circles.

(B) Magnified views of the yellow boxed regions in (A).

(C) Vascular density in the basolateral complex of AD and normal amygdala specimens.

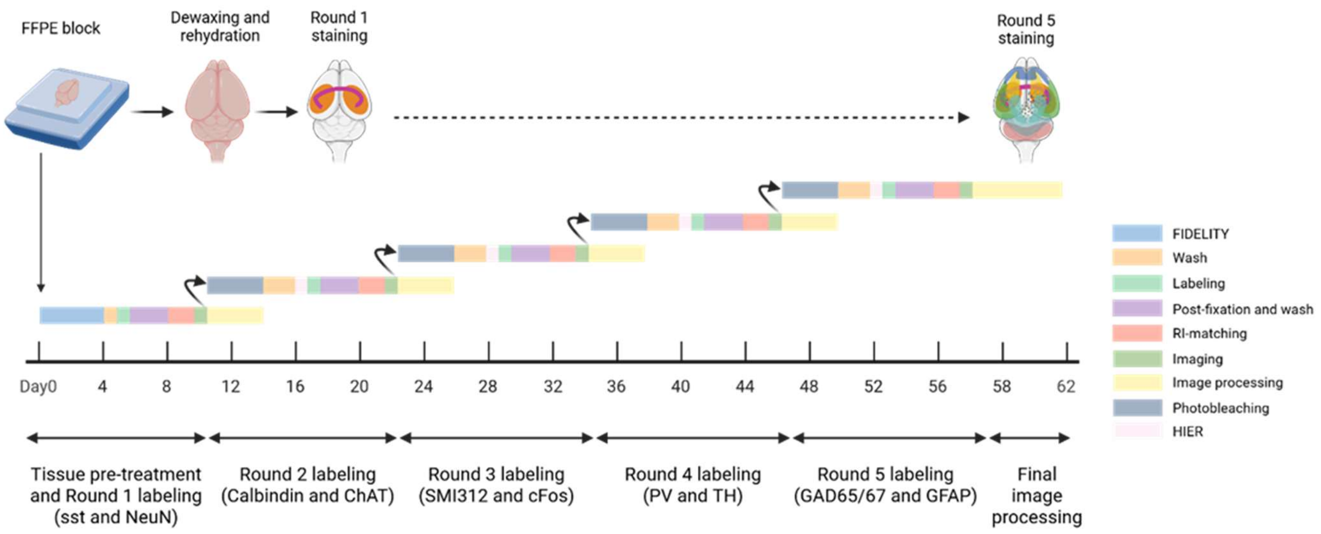


**Extended Data Figure 7. Timeline of five-round multiplexed whole-brain immunostaining using FIDELITY.**

Schematic timeline of the five-round multiplexed immunostaining workflow for whole mouse brains using FIDELITY, corresponding to Figures 4 and 5. (Created in https://BioRender.com)

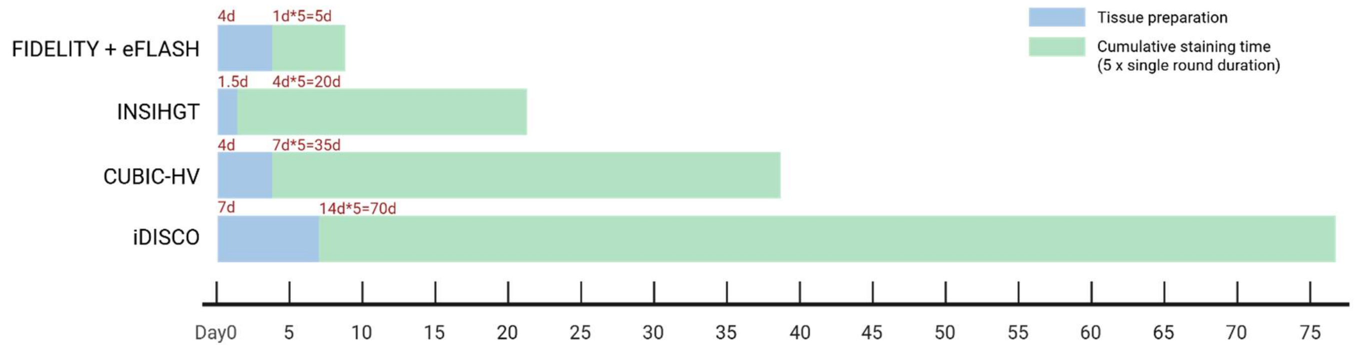


**Extended Data Figure 8. Comparison of tissue preparation and cumulative staining times for five single-round equivalents.** Protocol-based comparison of time requirements for whole-mouse-brain processing using FIDELITY + eFLASH, INSIHGT, CUBIC-HV, and iDISCO. Blue segments indicate one-time tissue preparation, counted from the start of fixation following tissue collection through completion of the preparatory steps preceding immunostaining. For FIDELITY, this interval includes fixation and paraffin-block preparation, rather than starting from a pre-existing FFPE block. Green segments indicate cumulative staining time, calculated by multiplying the duration of one staining round by five. Single-round staining includes antibody incubations and associated washing steps, with both primary- and secondary-antibody labeling included where applicable. Single-round staining durations were 1, 4, 7, and 14 days for FIDELITY + eFLASH, INSIHGT, CUBIC-HV, and iDISCO, respectively, corresponding to cumulative staining times of 5, 20, 35, and 70 days. Refractive index matching, imaging, and inter-round processing, including fluorescence inactivation or antibody elution and associated washes, were excluded. These estimates compare defined preparation and staining steps and do not represent the total duration or establish the feasibility of five-round cyclic workflows for all methods.

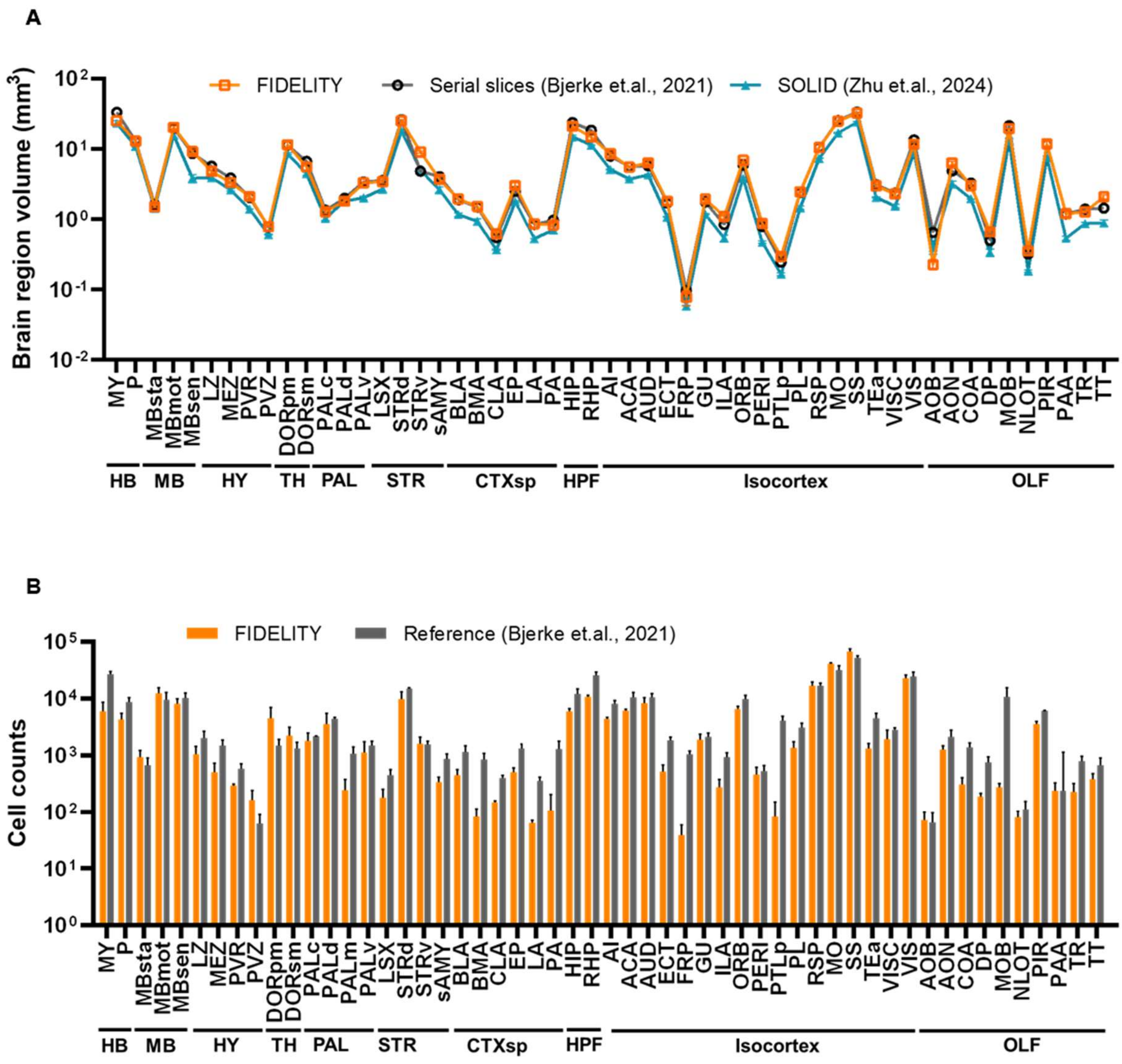


**Extended Data Figure 9. Comparison of brain region volumes (A) and parvalbumin interneuron cell numbers (B) computed from brains processed with other protocols.**

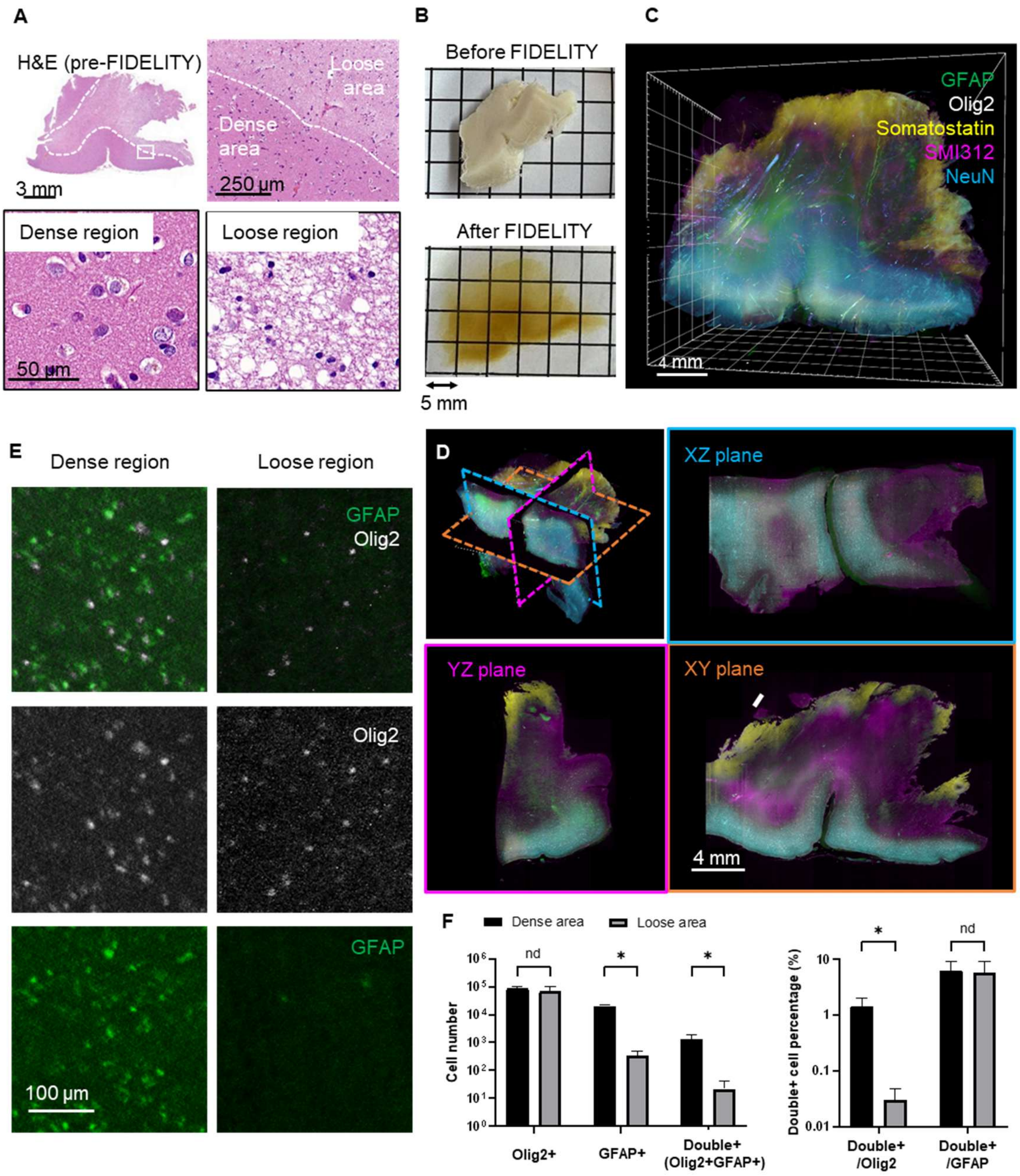


**Extended Data Figure 10. FIDELITY enables 3D cell phenotyping of a human glioma specimen.**

(A) Hematoxylin-and-eosin (H&E) images of the glioma specimen prior to FIDELITY processing.

(B) Macroscopic images of the same specimen before and after FIDELITY, illustrating tissue clearing.

(C) 3D rendering of a five-color image showing the entire specimen.

(D) Optical sections of the XZ, YZ, and XY planes.

(E) Magnified views showing the distribution of Olig2 and GFAP in dense and loose regions.

(F) Quantification of $Olig2^+$, $GFAP^+$, and $Olig2^+/GFAP^+$ double-positive cell populations in dense versus loose regions.

Statistical analysis: n = 5 ROIs per region; data represent mean ± SD.

nd: not significant ($P > 0.1$); $GFAP^+$ $P < 0.000001$; $Double^+$ $P = 0.0021$; $Double^+/Olig2^+$ $P = 0.0006$.

Multiple unpaired t-tests were performed.
